# GASTON-VI efficiently quantifies spatial axes and molecular gradients from multi-sample, multi-modal spatial-omics data

**DOI:** 10.64898/2026.11.24.769044

**Authors:** Zhirui Ning, Sayuni Dharmasena, Uthsav Chitra

## Abstract

Many tissues exhibit an underlying *spatial axis* along which fundamental biological processes occur due to continuous spatial gradients of cell type and molecular state. Modern spatial-omics technologies measure molecular features (e.g. gene/protein expression) at 100,000s-1,000,000s of locations within a two-dimensional (2-D) tissue slice, with large-scale atlases profiling multiple tissue slices of the same organ type across different individuals and biological conditions (e.g. healthy versus diseased). However, existing computational methods for identifying spatial molecular gradients are limited to small spatial transcriptomics datasets (1000s of cells) of a single tissue slice, while methods for integrating multiple samples do not explicitly model spatial axes nor spatial molecular gradients along such axes. Here, we present GASTON-VI, a variational inference (VI)-based deep learning model which efficiently learns an underlying spatial axis and spatial molecular gradients along this axis from multi-sample, multi-condition spatial-omics data of any molecular modality (e.g. transcriptomics, proteomics). GASTON-VI jointly learns (1) a spatial coordinate (the *“isodepth”*) for each individual tissue slice and (2) 1-D spatial modality functions that describe molecular gradients along the isodepth for tissues of the same biological condition, while utilizing a latent variable model to account for batch effects. Further, GASTON-VI implements a statistical test for identifying molecules with *differential* spatial gradients across biological conditions (e.g. healthy versus diseased). On spatial transcriptomics and proteomics data from the intestine, tonsil, and liver, we show that GASTON-VI recovers spatial axes and molecular gradients along these axes more accurately and efficiently than existing methods. In particular, applied to large-scale spatial transcriptomics and proteomics atlases, GASTON-VI reveals differential spatial gradients of gene expression along the portal vein-central vein axis in fibrotic livers, and differential spatial gradients of protein abundance along the muscularis-lumen axis across different regions of the intestine.

**Software availability:** GASTON-VI is available at https://github.com/chitra-lab/GASTON-VI.

## 1 Introduction

A central problem in spatial biology is to systematically characterize spatial variation in cellular and molecular state in biological tissues. Many tissues exhibit an underlying *spatial axis* along which fundamental biological processes occur, typically due to continuous spatial variation (or *spatial gradients*) in cell type composition and molecular cell state along this axis. For example, in the liver, hepatocyte division of labor is orchestrated through continuous gradations in metabolic cell state along the portal vein-central vein (PV-CV) axis [1, 2]; in the intestine, stem cell migration and differentiation occur along the crypt-tip axis of individual villi [3, 4]; and in tumors, spatial gradients of oxygen and metabolites along the stroma-tumor core axis promote angiogenesis and are a key biomarker of cancer progression [5, 6]. Characterizing the underlying spatial axis of a tissue and spatial molecular gradients along this axis is critical for understanding biological function and disease [7– 10].

Modern high-throughput spatial-omics technologies measure molecular features at thousands to millions of spatial locations in one or multiple two-dimensional (2-D) tissue slices. For example, spatial transcriptomics (ST) measures RNA transcripts (gene expression) inside individual cells [11–14], while spatial proteomics (SP) and spatial metabolomics (SM) measure protein and metabolite abundance, respectively [15–20]. However, these technologies provide sparse or incomplete molecular measurements; sequencing-based technologies measure a small number of unique molecular identifiers (UMIs) at each spatial location (approximately 100-500 UMIs per 10*µ*m ×10*µ*m region [21, 22]), while imaging-based technologies measure relatively small and targeted subsets of molecular features in a tissue (typically tens to thousands of features [23]). This sparsity markedly complicates the identification and interpretation of spatial molecular variation along underlying tissue axes from spatial-omics data.

There are relatively few approaches to identify spatial axes of molecular variation from spatial-omics data. A commonly used approach is to learn a low-dimensional embedding vector for each cell using approaches such as matrix factorization [24–27] or deep learning [28–31], and manually identify an embedding component that corresponds to a spatial axis. However, the low-dimensional embeddings may not be spatially coherent and thus may not have components that correspond to a biologically meaningful spatial axis. One of the authors recently developed GASTON, a deep learning model for learning a spatial axis called the *isodepth* from ST data and parametrizing both continuous gene expression gradients and discontinuities in expression across this axis [32]. However, GASTON’s relatively simple neural network architecture limits its expressivity, making it unable to learn spatial axes that repeat across anatomical units of a tissue, such as the PV-CV axis that repeats across liver lobules [2], nor scale to large datasets with hundreds of thousands to millions of cells. Furthermore, GASTON does not model non-transcriptomics molecular modalities such as spatial proteomics.

Many recent studies generate large *spatial-omics atlases* that profile 2-D tissue slices from the same organ type across different individuals and biological conditions (e.g., healthy versus diseased). In the field of single-cell omics, many computational methods have been developed to integrate multiple single cell datasets while accounting for *batch effects*, or technical variation across datasets (e.g. see reviews/benchmarks in [33–35]), including the widely-used *variational inference (VI)*-based methods such as scVI [36]; some recent methods now integrate multiple spatial datasets [37–50]. However, these methods do not explicitly model spatial axes of molecular variation and therefore are unable to identify spatial molecular gradients from multi-sample spatial-omics data, nor identify *differential* spatial gradients across different biological conditions. On the other hand, methods for identifying spatial axes of variation such as GASTON are only applicable to an individual tissue slice. Thus, to our knowledge, there are no existing computational methods that identify spatial axes and molecular gradients along such axes from multiple tissue slices from different individuals and biological conditions.

Here, we present GASTON-VI, a deep learning model that efficiently learns an underlying spatial axis and spatial molecular gradients along this axis from multi-sample, multi-condition spatial-omics data of any molecular modality (e.g. transcriptomics, proteomics). Given spatial-omics data of multiple tissue slices, GASTON-VI jointly learns an isodepth coordinate for each tissue slice and one-dimensional (1-D) spatial modality functions that describe molecular variation (spatial gradients) along the isodepth. GASTON-VI leverages several methodological improvements to scale to large spatial-omics data of any molecular modality: (1) a neural field model that uses Fourier feature positional encodings [51] to learn more expressive isodepth coordinates compared to the original GASTON, and (2) a VI-based framework that accounts for batch effects by learning cell state latent vectors and (3) a modality-specific probabilistic model to account for different molecular modalities. Further, GASTON-VI enables *differential* spatial gradient analysis by learning 1-D spatial modality functions that are shared by tissues of the same biological condition but may differ across conditions, e.g. demonstrating how spatial molecular gradients vary across tissues from healthy versus diseased individuals. We apply GASTON-VI to spatial-omics data across a range of diverse organ types (intestine, tonsil, liver), modalities (transcriptomics, proteomics), and biological conditions (healthy versus diseased, intestinal sections) and show that GASTON-VI more efficiently and accurately recovers underlying spatial axes and molecular gradients along these axes than existing methods.

## 2 Results

### 2.1 GASTON-VI learns spatial axes and spatial molecular gradients from multi-sample spatial omics data

We introduce GASTON-VI, a method for learning the underlying spatial axis(es) of variation and spatial molecular gradients along these axes from spatial-omics of one or more tissue slices and one or more biological conditions (e.g. healthy, diseased; Fig. 1). GASTON-VI has the following features: (1) for each tissue slice, GASTON-VI learns a 1-D spatial axis called the *isodepth* which describes the axis of maximum spatial variation in molecular abundance [32, 52, 53], and *spatial gradients* of molecular abundance along the isodepth; (2) GASTON-VI uses a variational inference (VI)-based framework [36, 54] to efficiently learn the isodepth and spatial gradients from large-scale data; (3) GASTON-VI is applicable to multiple molecular modalities including transcriptomics (e.g. MERFISH [55], 10x Genomics Visium HD [56] and Xenium [57]) and proteomics (e.g. CODEX [15]); and (4) GASTON-VI implements a statistical test to identify *differential spatial gradients* across tissues from different biological conditions (e.g., healthy versus diseased).

**Figure 1:**
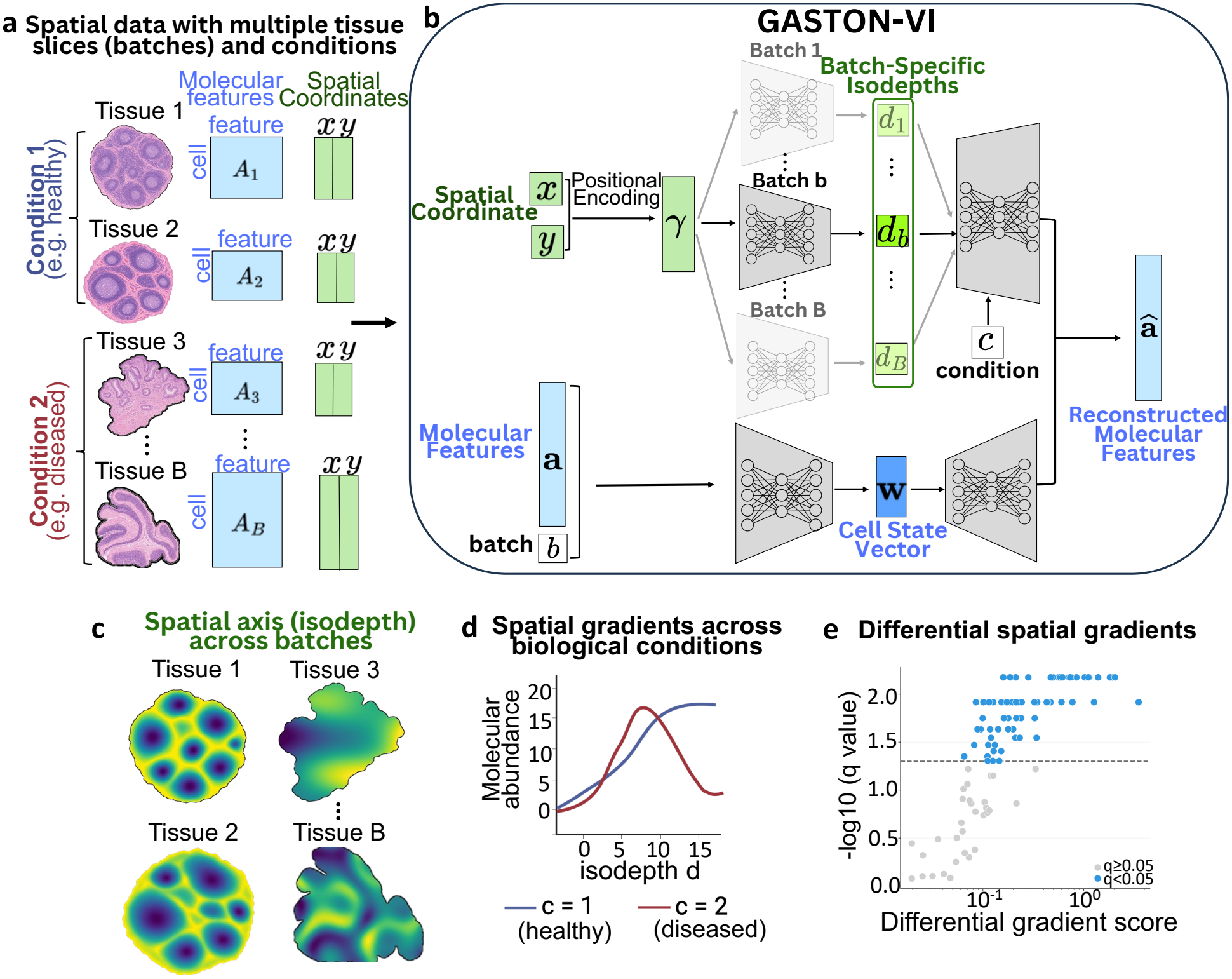
**a**, GASTON-VI takes as input spatial-omics data from multiple tissue slices across different biological conditions. Each tissue slice is treated as a batch *b* ∈ {1, …, *B*}, with a molecular feature matrix *A*_*b*_, a spatial coordinate matrix *S*_*b*_ containing the spatial coordinates (*x, y*) of its cells, a discrete batch label *b*, and a discrete condition label *c*_*b*_. **b**, GASTON-VI is a variational inference (VI)-based deep learning model which takes as input spatial coordinates (*x, y*), molecular feature vector **a**, batch label *b*, and condition label *c* of a cell and outputs a reconstructed molecular feature *â*. GASTON-VI applies a Fourier feature positional encoding *γ* = *γ*(**s**) to the spatial coordinates **s** and uses one of *B* neural networks to compute a *batch-specific isodepth d*_*b*_; simultaneously, GASTON-VI passes the molecular features **a** and batch label *b* into a neural network to compute a *cell state vector* **w.** GASTON-VI then reconstructs the molecular feature vector **â** using *h*_*m*_(*d*_*b*_, *c*) + *e*_*m,b*_(**w**), given the batch-specific isodepth *d*_*b*_, condition label *c*, and cell state vector **w**, where the functions *h*_*m*_, *e*_*m,b*_ are neural networks. GASTON-VI learns **c**, an isodepth coordinate *d*_*b*_ for each batch *b* = 1, …, *B*, which models the underlying spatial axis shared across batches, and learns **d**, a scalar function *h*_*m*_(*d, c*) which describes how molecular feature *m* varies along the isodepth *d*, i.e. *spatial molecular gradients*. **e**, GASTON-VI implements an efficient statistical test to identify molecular features *m* with significant differential spatial gradients between conditions *c, c*′, i.e. *h*_*m*_(*d, c*) ≠ *h*_*m*_(*d, c*′).

Briefly, GASTON-VI takes as input (Fig. 1a) for each tissue slice (batch) *b*: a molecular feature matrix *A*_*b*_ (e.g. gene expression matrix), a spatial coordinate matrix **S**_*b*_ where each row **s** = (*x, y*) is the 2-D spatial coordinates for each cell, and (optionally) a discrete condition label *c*_*b*_, such as the disease/developmental state or the organ of slice. For each tissue slice *b* with biological condition *c*, GASTON-VI models the abundance *f*_*m,b*_ of molecular feature *m* in a cell as the following additive function of (1) an isodepth *d*_*b*_(**s**), or a one-dimensional (1-D) coordinate describing the relative position of spatial location **s** on the underlying axis of the tissue slice; and (2) a low-dimensional cell state vector **w**:

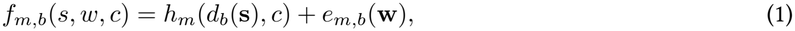

where *h*_*m*_(*d*_*b*_, *c*) is a scalar function that describes *spatial molecular gradients*, or how molecular abundance varies along the shared isodepth *d*_*b*_, and *e*_*m,b*_(**w**) is a scalar function that describes the contribution of cell state **w** to molecular abundance (Methods).

GASTON-VI uses deep learning-based variational inference (VI) to efficiently learn the tissue slice-specific isodepth functions *d*_*b*_, spatial gradient functions *h*_*m*_, cell state vectors **w**, and cell state functions *e*_*m,b*_ (Figure 1b). GASTON-VI uses modality-specific probabilistic likelihoods to model different molecular modalities, and incorporates Fourier feature positional encoding [51] and an efficient, two-stage training scheme (Methods).

A key distinguishing feature of GASTON-VI is its interpretable model of molecular abundance (Eq. (1)) which helps overcome the substantial data sparsity of current spatial-omics technologies. Specifically, by modeling spatial molecular abundance in terms of a 1-D isodepth coordinate *d*_*b*_(**s**) (Figure 1c), GASTON-VI implicitly aggregates information across different locations **s** with the same isodepth *d*_*b*_. Furthermore, by sharing the spatial gradient functions *h*_*m*_ across different tissue slices *b* of the same condition *c* (Figure 1d), GASTON-VI aggregates information across multiple tissue slices. Another important feature of GASTON-VI is that it implicitly disentangles spatial and non-spatial molecular variation by separating the contributions of spatial molecular gradients (through the functions *h*_*m*_) and molecular cell state (through the functions *e*_*m,b*_) to the overall molecular abundance of each cell.

Another core contribution of GASTON-VI is its ability to identify *differential* spatial gradients across biological conditions (Figure 1e). Specifically, GASTON-VI implements a highly efficient statistical test using a parametric bootstrap to determine whether two biological conditions *c, c*′ (e.g. healthy, diseased) have different spatial gradient functions *h*_*m*_(·, *c*), *h*_*m*_(·, *c*′), respectively. GASTON-VI quantifies these differences using a differential gradient (DG) score *D*_*m*_, where higher scores indicate larger differences between the spatial gradients *h*_*m*_(·, *c*), *h*_*m*_(·, *c*′) across conditions *c, c*′. GASTON-VI’s efficient statistical test enables the identification of novel differential spatial gradient patterns in atlas-scale spatial-omics data.

### 2.2 GASTON-VI recapitulates known spatial zonation gradients in mouse liver

We first used GASTON-VI to identify spatial gradients of gene expression in a MERFISH mouse liver ST dataset, which measures the expression of 347 genes across 367, 335 cells in a single tissue slice [58]. The liver is comprised of repeating hexagonal structures called lobules. Several studies have found that many liver-related genes exhibit continuous *spatial gradients* of gene expression along a portal vein-central vein (PV-CV) axis inside each lobule, a phenomenon known as *liver zonation* [1, 2, *59*] *(Fig. 2a, left). Such genes are typically classified as either periportal genes* or *pericentral genes*, depending on whether their expression decreases or increases along the PV-CV axis (see Fig. 2a, right). However, current computational methods struggle to infer both the PV-CV axis and spatial gradients along the PV-CV axis from large-scale ST data. For example, our previous method GASTON could only identify the PV-CV axis and spatial gradients from small, manually cropped regions containing individual lobules, a time-consuming and potentially error-prone process [32, 53]. We hypothesized that GASTON-VI’s improved architecture would allow us to efficiently infer the PV-CV axis and spatial gradients across the entire dataset.

**Figure 2:**
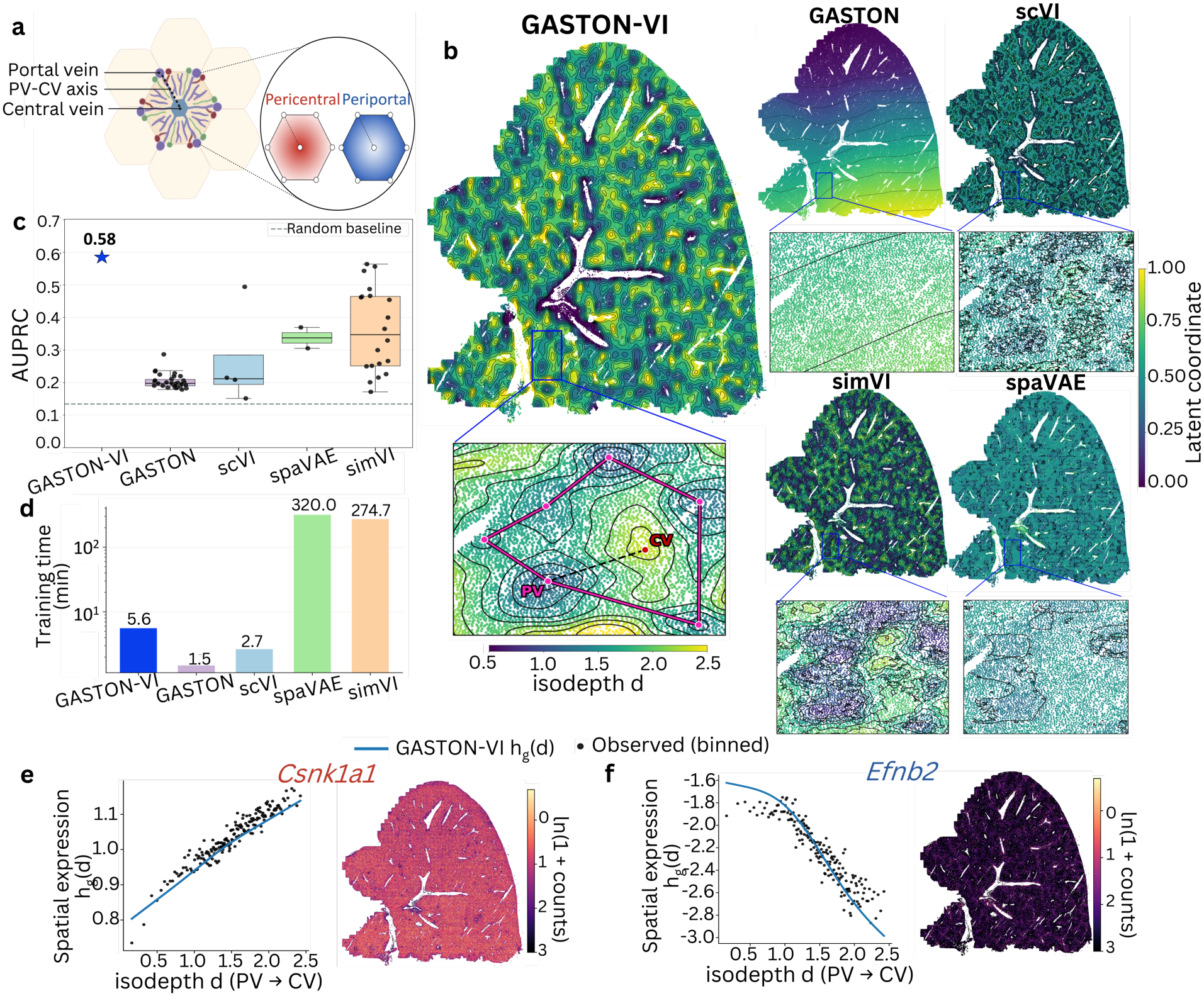
**a**, Schematic of repeating liver lobules and spatial expression gradient of pericentral and periportal genes within lobules, with darker shading indicating higher expression. **b**, GASTON-VI isodepth and the latent coordinate inferred by comparison methods. Black curves denote contours of equal latent values. Insets show the region corresponding to a single hepatic lobule. In the GASTON-VI inset, the central vein (CV) corresponds to a local isodepth maximum, and the portal veins (PVs) correspond to local isodepth minima. **c**, Recovery of known liver zonation markers measured by macro-averaged directional area under the precision-recall curve (AUPRC). For comparison methods, black dots show AUPRC values from individual runs or latent dimensions. **d**, Training times for GASTON-VI and comparison methods. **e-f**, GASTON-VI inferred 1-D spatial gene expression functions *h*_*g*_(*d*) (left) and observed 2-D expression plots (right) for *Csnk1a1* (**e**) and *Efnb2* (**f**). Black dots denote the observed counts transformed to the same scale as spatial expression functions *h*_*g*_.

The isodepth *d* learned by GASTON-VI visually matches the 1-D PV-CV axis in the liver (Fig. 2b). Within hepatic lobules, the GASTON-VI isodepth peaks at the central vein and decreases continuously toward the surrounding portal veins, thus capturing successive positions along the 1-D PV-CV axis. To quantitatively evaluate our learned isodepth *d*, we benchmarked GASTON-VI against 1-D spatial coordinates learned by four other methods, evaluating how well such learned coordinates identify known liver zonation genes. The comparison methods include simVI [28] and spaVAE [29], spatial latent variable models trained using variational inference; scVI, a widely used single-cell latent variable model trained using variational inference [36]; and GASTON, our model’s predecessor. For each method, we fit the expression *a*_*ig*_ of each gene *g* against the learned coordinate using linear regression and ranked genes by their slopes *β*, where ideally a large positive slope (*β≫* 0) corresponds to pericentral markers and a large negative slope (*β ≪* 0) corresponds to periportal markers. We evaluated the rankings against known pericentral and periportal markers [1, 2, 59–61] using macro-averaged area under the precision-recall curve (AUPRC) in a one-vs-rest (OvR) setup; see Method 4.6 for details.

GASTON-VI more accurately identifies the PV-CV axis and known liver zonation genes than the other methods (Figure 2b-c). Visually, the GASTON-VI isodepth is the only coordinate which both smoothly varies across the tissue slice and captures the repeating lobule geometry. On the other hand, the GASTON isodepth does not repeat across the tissue due to its simpler deep learning architecture, while the coordinates learned by scVI, spaVAE, and simVI do not smoothly vary within lobules as these methods do not explicitly model smooth spatial variation (Figure 2b). By learning a more spatially coherent PV-CV axis, GASTON-VI more accurately identifies known liver zonation genes than existing methods (Figure 2c; GASTON-VI AUPRC = 0.58 versus median AUPRC *<* 0.35 for other methods). We observed that the AUPRC of scVI, spaVAE, and simVI varied considerably across the different coordinates learned by each method, highlighting the difficulty of identifying an interpretable spatial axis from a large list of learned latent variables. In contrast, GASTON-VI learns a single, interpretable spatial coordinate—the isodepth *d*—that describes the spatial axis of maximum variation in gene expression.

We emphasize that GASTON-VI is highly efficient to train, taking only 5.6 minutes to train on this MERFISH dataset with over 300, 000 cells (Fig. 2d). The GASTON-VI runtime is comparable to GASTON and scVI, and is substantially faster than the spatial latent variable models spaVAE and simVI which require over 4 hours.

GASTON-VI reveals spatial expression gradients that are not readily apparent in the raw data by learning a continuous 1-D spatial gene expression function *h*_*g*_(*d*) for each gene that models how gene expression varies along the isodepth *d*. For example, the spatial expression function *h*_*g*_(*d*) for the gene *Csnk1a1* increases smoothly along the isodepth from the portal vein towards the central vein (Fig. 2e, left), consistent with the classification of *Csnk1a1* as a pericentral marker [1, 2, 59, 61]. However, such a spatial gradient is difficult to observe when visualizing the raw expression values in 2-D (Fig. 2e, right). Similarly, the GASTON-VI spatial expression function *h*_*g*_(*d*) for the gene *Efnb2* decreases smoothly along the isodepth from the portal vein towards the central vein (Fig. 2f, left), which is consistent with its classification as a periportal marker [1, 2, 59, 61]; this spatial expression gradient is less readily apparent when visualized in 2-D (Fig. 2f, right). As further validation of the GASTON-VI-learned spatial gradients *h*_*g*_(*d*), we used GASTON-VI to jointly analyze four MERFISH liver tissue samples from two healthy mice and found that the spatial expression functions *h*_*g*_(*d*) for known periportal and pericentral genes are consistent between the single-sample and multi-sample analyses; see Appendix S3 for details.

One reason that GASTON-VI accurately recovers spatial gene expression gradients along the PV-CV axis is that GASTON-VI separately models cell-intrinsic variation through a learned cell state vector **w**. To demonstrate the importance of the cell state vector, we show that if we use a variant of GASTON-VI that does not learn a cell state vector **w**, then GASTON-VI learns spatial expression functions *h*_*g*_(*d*) that less accurately reflect the known liver zonation patterns (Appendix S4).

### 2.3 GASTON-VI reveals differential spatial gradients between healthy and fibrotic livers

We next evaluated GASTON-VI using a large-scale MERFISH ST atlas, which consists of three batches (tissue slices) from healthy human livers and five batches from fibrotic human livers, measuring the expression of *G* = 317 genes across 575, 666 cells [62] (Fig. 3a). GASTON-VI learns batch-specific isodepths *d*_*b*_ that visually recover the PV-CV axis in both healthy and fibrotic livers (Fig. 3b). Interestingly, the isodepth range is noticeably smaller for fibrotic livers than for healthy livers (Fig. 3c), suggesting a change in the PV-CV spatial organization in fibrosis.

**Figure 3:**
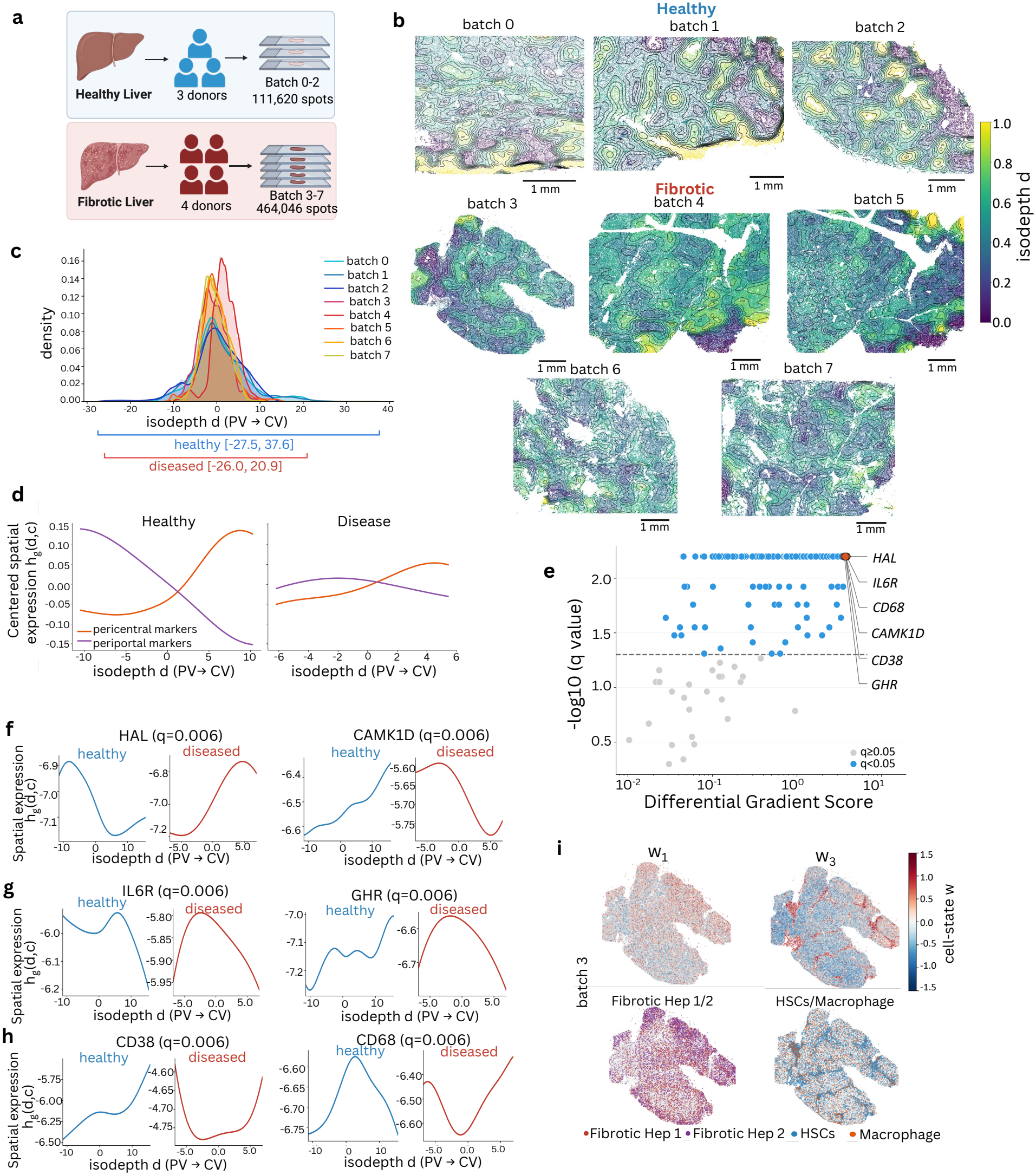
**a**, Schematic of human MERFISH liver dataset [62]. **b**, GASTON-VI isodepths *d*_*b*_ learned across healthy and fibrotic liver samples *b*. Isodepths *d*_*b*_ are min-max normalized across all tissue slices to lie in the range [0, 1]. **c**, Distribution of the inferred isodepth for each batch. **d**, Mean-centered spatial gene expression functions *h*_*g*_(*d, c*) averaged across known pericentral and periportal marker genes *g* across healthy and fibrotic conditions *c*. **e**, Scatterplot of GASTON-VI-estimated differential spatial gradient score versus −log_10_(*q*) across different genes measured in the human MERFISH liver dataset. Each dot is a gene, with a dashed line drawn at *q* = 0.05 and blue dots indicating genes with *q* < 0.05. **f-h**, Spatial expression functions of six genes with significant differential spatial gradients between healthy and fibrotic conditions. **i**, (Top) GASTON-VI cell state vector *w*_1_ and *w*_3_ for fibrotic liver batch 3; (bottom) cell type annotations from the original study.

The spatial expression functions *h*_*g*_(*d, c*) learned by GASTON-VI reveal different liver zonation patterns between healthy and fibrotic conditions, including an *attenuation* (decrease) in hepatocyte zonation in fibrotic livers. In both the healthy and fibrotic conditions *c*, the learned spatial expression functions *h*_*g*_(*d, c*)—averaged over known peri-central and periportal marker genes *g*—follow the expected increasing and decreasing spatial patterns, respectively, along the PV-CV axis (Fig. 3d). However, the range of the spatial expression functions *h*_*g*_(*d, c*) is noticeably smaller for the fibrotic condition compared to the healthy condition (Fig. 3d, right), indicating a substantial *attenuation* of liver zonation in fibrotic livers. To demonstrate that such attenuation is real, we performed a negative control experiment where we randomly split cells in the healthy batches into two synthetic conditions and found negligible differences between the learned spatial expression functions (Supplementary Fig. S3b-c). The observed attenuation of zonation gradients for pericentral and periportal markers in fibrotic livers is consistent with previous observations of weakened hepatocyte zonation in diseased liver [63] and periportalization in liver fibrosis, where pericentral regions lose their pericentral identity and acquire periportal gene expression profiles [64].

A key feature of GASTON-VI is its ability to systematically determine which genes have *differential* spatial expression gradients across healthy and fibrotic livers. Briefly, GASTON-VI computes a differential gradient score (DG score) *D*_*g*_, which describes the difference (normalized mean squared distance) between the spatial gradients in the healthy and diseased livers, and implements a statistical test (parametric bootstrapping) to determine which genes *g* have a significantly larger DG score *D*_*g*_ than expected by chance. GASTON-VI identifies 239/317 genes with statistically significant differences (GASTON-VI *q <* 0.05) in spatial gradients between healthy and fibrotic livers (Figure 3e).

The genes that GASTON-VI identifies as having the most significant differential gradients (i.e. smallest *q* value) reveal altered zonation patterns of metabolism, regeneration, and immune response along the PV-CV axis. For example, the genes *HAL* (GASTON-VI *q* = 0.006, DG score *D* = 3.89) and *CAMK1D* (GASTON-VI *q* = 0.006, DG score *D* = 3.77) switch from exhibiting periportal zonation to pericentral zonation and vice versa, respectively (Fig. 3f).

*HAL* and *CAMK1D* are involved in metabolism in healthy livers [65, 66], indicating potential metabolism-related changes in fibrotic livers. Two other genes with the smallest *q*-values, *IL6R* (GASTON-VI *q* = 0.006, DG score *D*= 3.88) and *GHR* (GASTON-VI *q* = 0.006, DG score *D*= 3.64), were also identified by GASTON-VI as shifting from periportally and pericentrally zonated, respectively, to mid-zonated expression gradients in fibrotic livers (Fig. 3g), where their expressions peak at intermediate isodepths *d. IL6R* and *GHR* are involved in liver regeneration and repair [67–69], so the differential spatial gradient identified by GASTON-VI is consistent with the mid-zonal hepatocytes being the major cellular source for hepatocyte regrowth following injury [70]. GASTON-VI also identified two immune signaling genes *CD68* (GASTON-VI *q* = 0.006, DG score *D*= 3.78) and *CD38* (GASTON-VI *q* = 0.006, DG score *D*= 3.75) [71, 72] as having the most significant change in spatial gradients, with *CD68* and *CD38* shifting from mid-zonated and pericentrally zonated, respectively, to mid-zonated expression gradients with small expression at intermediate isodepths *d* in fibrotic livers (Fig. 3h). This finding suggests an altered zonation of immune response along the PV-CV axis. Furthermore, by clustering the distinct spatial expression gradients *h*_*g*_(*d, c*) within each condition *c*, we identified five clusters of spatial gradients in healthy livers (Supplementary Fig. S3d) and five such clusters in fibrotic livers (Supplementary Fig. S3e) which visually correspond to the zonation patterns we highlight in Fig. 3f-h. See Appendix S2.7 for details.

Another key contribution of GASTON-VI is the learned cell state vector **w**, which models distinct cell populations in fibrotic livers. For example, the first and third entries of the learned cell state vector **w** capture cell populations annotated in the original study [62]: the first entry *w*_1_ corresponds to the fibrotic hepatocyte population, which is uniquely present in fibrotic livers and does not exhibit a clear zonation gradient along the PV-CV axis, while the third entry *w*_3_ captures the hepatic stellate cell (HSC) and macrophage populations (Fig. 3i).

Together, our results demonstrate that GASTON-VI efficiently identifies differential spatial gradients and cell populations from large-scale spatial-omics data of healthy and fibrotic livers, and indicate several potential biomarkers for liver fibrosis.

### 2.4 GASTON-VI identifies epithelium-TCZ-follicle axis and spatial expression gradients in human tonsil

We next applied GASTON-VI to a 10x Genomics Visium HD ST sample of the human tonsil [73], which measures the expression of *G =* 18, 085 genes across *N =* 553, 820 spatial locations. Human tonsils consist of repeating, B-cell-rich follicles separated by interfollicular T-cell zones (TCZs) along with surface and crypt epithelium [74, 75]. Prior biological studies have identified a few gene expression gradients within follicles as well as spatially localized gene expression patterns in the interfollicular regions [75, 76]. Since this Visium HD dataset of the tonsil measures the expression of all genes across the whole transcriptome in a large tissue slice, we hypothesized that GASTON-VI would be able to more systematically identify gene expression gradients within and between follicles in the tonsil. GASTON-VI learns a continuous isodepth coordinate *d* that increases from the epithelium through the interfollicular TCZs and within the follicles (Fig. 4a,b). In contrast, the isodepth learned by GASTON does not capture the repeating nature of follicles due to its simpler architecture, while the latent coordinates learned by scVI, spaVAE, and simVI do not smoothly vary within follicles as these methods do not explicitly model smooth spatial variation (Fig. 4c). Furthermore, GASTON-VI is highly efficient, taking only 14 minutes to train. GASTON-VI has a comparable runtime to GASTON and scVI while being 7-12 × faster than simVI and spaVAE, which are generative models explicitly designed for identifying spatial patterns (Fig. 4d).

**Figure 4:**
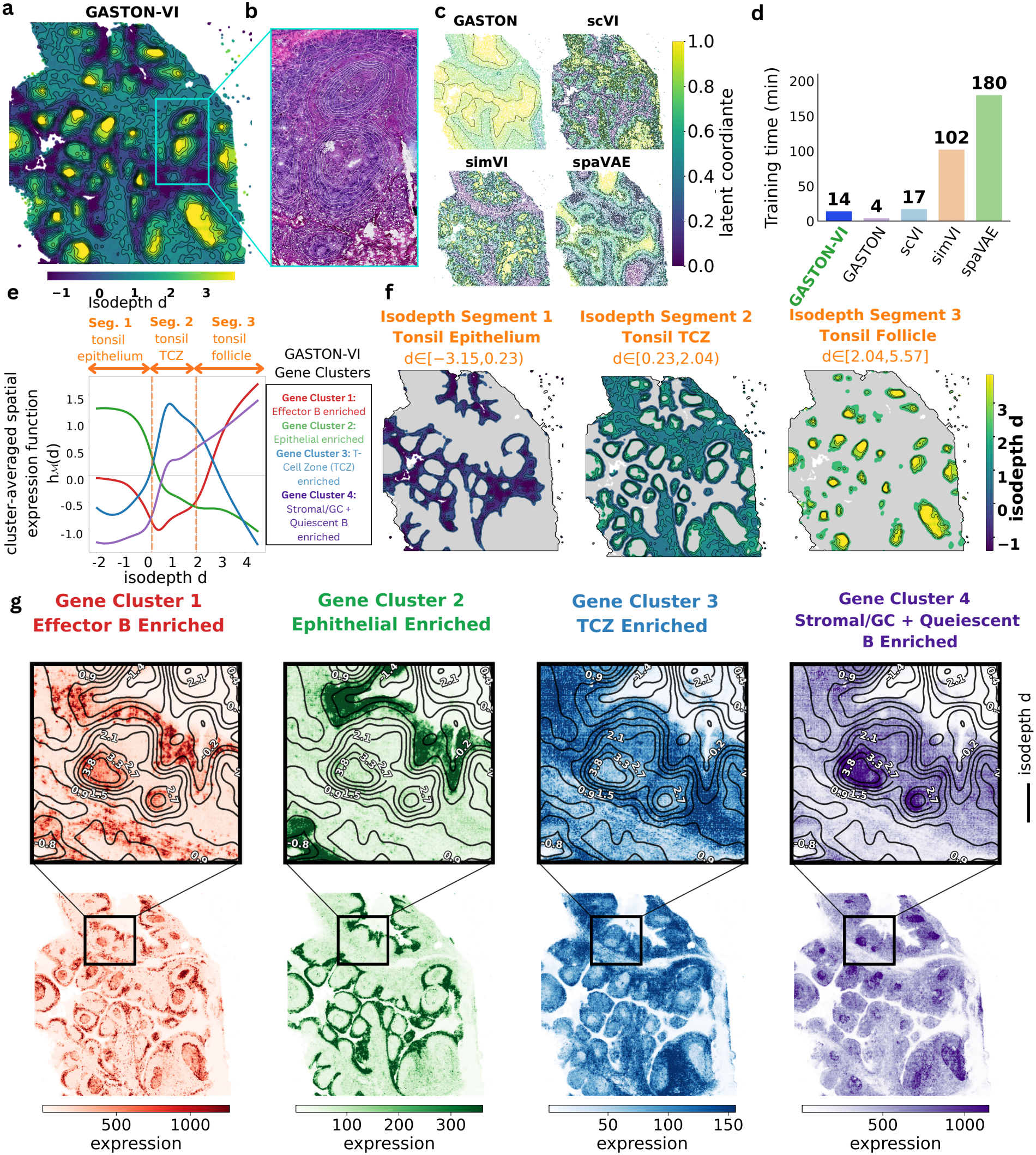
**a**, GASTON-VI isodepth *d* inferred from the Visium HD human tonsil dataset [73]. Black contour curves denote equal isodepth values. **b**, H&E image of the boxed region in **a. c**, Latent coordinates inferred by comparison methods GASTON, scVI, simVI, and spaVAE. **d**, Training times for GASTON-VI and comparison methods. **e**, Cluster-averaged spatial expression functions *h*_*ℳ*_ *(d)* for four gene clusters identified by *K*-means clustering of the spatial expression functions *h*_*g*_ (d). Clusters are labeled according to enriched pathways from GSEA. Orange dashed vertical lines denote breakpoints identified by Belayer [77]. Orange arrows mark the three isodepth segments mapped in **f**, corresponding to the tonsil epithelium (segment 1), T-cell zone (segment 2), and follicle (segment 3). **f**, Isodepth *d* restricted to each of the three segments shown in **e.** Black contour curves denote equal isodepth values. **g**, Observed 2-D spatial expression aggregated across genes in each of the four clusters shown in **e**. Top panels show enlarged views of the boxed regions in the tissue maps below. Black contour curves denote equal isodepth values.

The GASTON-VI isodepth *d* describes an epithelium-TCZ-follicle axis that underlies the tonsil. We validated this interpretation of the isodepth in two ways. First, we performed a piecewise linear regression (implemented in [77]) jointly across the learned spatial expression functions *h*_g_ (*d*), which partitioned the isodepth into three segments (Fig. 4e). By performing a gene set enrichment analysis (GSEA) [78] on the highly expressed genes in each segment, we found that segment 1 was enriched for epithelial cell pathways, segment 2 for naive T-cell pathways, and segment 3 for germinal center B-cell pathways (Supplementary Fig. S7a), indicating that the three segments correspond to epithelium, TCZ, and follicle regions, respectively (Fig. 4f). Second, to identify specific spatial gradients along the isodepth, we clustered the spatial expression functions *h*_g_ (*d*) learned by GASTON-VI to obtain four gene clusters (Fig. 4e; Methods). By performing a GSEA on each cluster, we found that cluster 1 was enriched for effector B-cell pathways, cluster 2 for epithelial pathways, cluster 3 for TCZ pathways, and cluster 4 for immune/stromal pathways (Supplementary Fig. S7b), which matches our interpretation of the GASTON-VI isodepth as modeling an epithelium-TCZ-follicle spatial axis.

GASTON-VI learns cluster-averaged spatial gradient functions *h*_*ℳ*_ (*d*) for different clusters *ℳ* of genes which reveal spatial gradients inside the tonsil. For example, GASTON-VI gene cluster 1 (effector B, Fig. 4e) exhibits an increasing spatial expression gradient within the third segment of the isodepth, which corresponds to tonsil follicles (Fig. 4f). GASTON-VI gene cluster 1 has a spatial gradient function 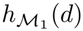 showing increasing expression toward the follicular interior, consistent with the enrichment of proliferating B cells in follicle germinal centers [79]. Such a smooth spatial gradient is less apparent when visualizing the sparse cluster-summed raw gene expression in 2-D (Fig. 4g). Similarly, the cluster-averaged expression functions *h*_*ℳ*_ (*d*) for GASTON-VI gene clusters 2 and *3* indicate spatial gradients of epithelial- and TCZ-related expression, with maximum expression in the epithelial and TCZ isodepth segments *d*, respectively (Fig. 4e,f) which matches our expectations [75]. Interestingly, GASTON-VI identifies a fourth gene cluster enriched for immune/stromal pathways (Fig. 4e) which has a sharp change in expression at the epithelium-TCZ boundary and a smoother spatial gradient in the TCZ and follicle. The spatial expression gradient of GASTON-VI gene cluster 4 may reflect the gradual increase of germinal center B-cells from the T-cell zone into B-cell-rich follicles [75]. In this way, GASTON-VI enables the identification of the epithelium-TCZ-follicle axis underlying the tonsil as well as subtle spatial expression gradients within follicles.

### 2.5 GASTON-VI recovers the muscularis-lumen axis and differential spatial protein gradients in a multi-patient human intestine dataset

We next applied GASTON-VI to a large-scale human intestine CODEX spatial proteomics atlas which measures the abundance of *M =* 49 proteins across *N =*2, 603, 217 cells in *B =* 64 tissue slices [80]. The tissue slices are collected from eight patients, with one tissue slice from each of eight different anatomical intestinal regions (duodenum, proximal jejunum, mid jejunum, ileum, ascending colon, transverse colon, descending colon, and sigmoid colon) per patient (Fig. 5a,b). We treated each tissue slice as a batch *b* and each intestinal region as a condition *c* [80]. The human intestine is organized along a muscularis-lumen axis and is known to exhibit gradients of immune cells along this axis [80] (Fig. 5c). Thus, we hypothesized that GASTON-VI’s general model of spatial axes would enable the identification of the muscularis-lumen axis across the different batches; spatial gradients of protein abundance along the muscularis-lumen axis; and whether any proteins exhibit differential spatial gradients across different intestinal regions.

**Figure 5:**
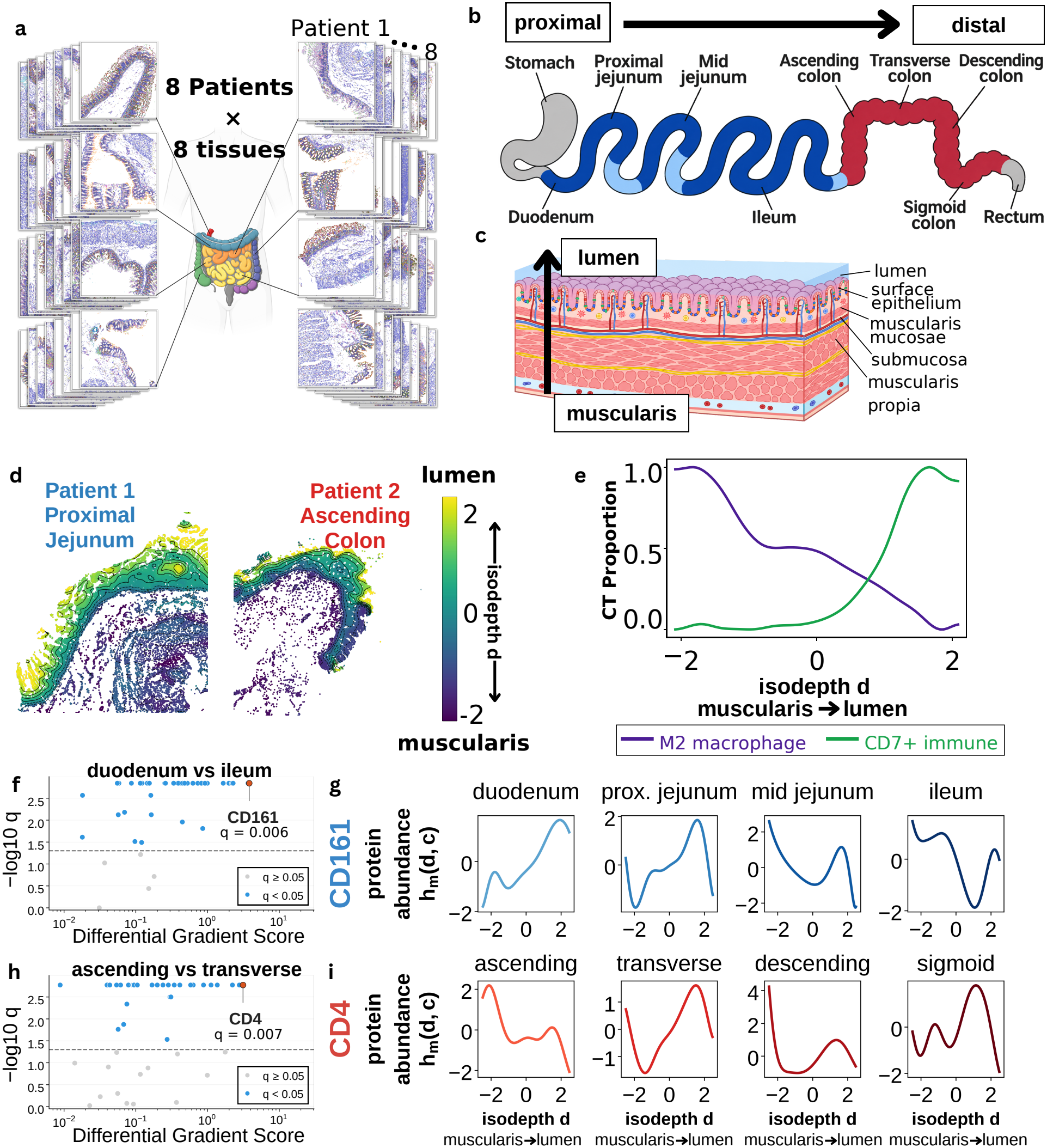
**a**, Schematic of the CODEX spatial proteomics dataset [80], showing tissue slices from eight intestinal regions in each of eight patients. **b**, Schematic of the human intestinal tract showing the eight sampled regions in proximal-to-distal order, from the duodenum to the descending sigmoid colon. **c**, Schematic of the intestinal wall showing the axis from the muscularis propria through the submucosa and mucosa to the lumen. **d**, GASTON-VI isodepth *d* in two representative tissue slices from the proximal jejunum of patient 1 (left) and the ascending colon of patient 2 (right). Cells are colored by inferred isodepth, from low values near the muscularis (purple) to high values near the lumen (yellow). Black contours connect locations of equal isodepth. **e**, Proportions of M2 macrophages (purple) and CD7^+^ immune cells (green) as functions of isodepth *d*, averaged across tissue slices. **f**, Differential gradient score versus – log_10_ (q) for each protein in the duodenum versus ileum comparison. CD161 is highlighted (q = 0.006). The dashed line marks *q* = 0.05. **g**, Inferred spatial abundance functions *h*_*m*_ *(d, c)* for CD161 in the duodenum, proximal jejunum, mid jejunum, and ileum. Isodepth increases from the muscularis to the lumen in each plot. **h**, Differential gradient score versus – log_10_*(q*) for each protein in the ascending versus transverse colon comparison. CD4 is highlighted (*q* = 0.007). The dashed line marks *q* = 0.05. **i**, Inferred spatial abundance functions *h*_*m*_ *(d, c)* for CD4 in the ascending, transverse, descending, and descending sigmoid colon, plotted from the muscularis to the lumen.

GASTON-VI learned an isodepth *d*_*b*_ for each batch (tissue slice) b that visually corresponds to the muscularis-lumen axis, with isodepth values increasing from the muscularis toward the lumen (Fig. 5d; Supplementary Fig. S4). We validated the inferred isodepths *d*_*b*_ using the cell type annotations provided in the original publication. The original study [80] reported that M2 macrophage proportions were highest near the muscularis and decreased toward the lumen, whereas CD7^+^ immune cell proportions were near zero at the muscularis side and increased toward the lumen (Fig. 3h of [80]). Consistent with their observations, we find that the average M2 macrophage proportion over all batches decreased with isodepth while the average CD7^+^ immune cell proportion increased with isodepth, where we calculate the average cell type proportion across all batches (Fig. 5e). Looking at individual batches, we observe that M2 macrophage proportion decreased with isodepth in 60 of 64 tissue slices and CD7^+^ immune cell proportions increased in 45 of the 57 slices with sufficient CD7^+^ immune cells (Supplementary Fig. S6). Our findings indicate that the GASTON-VI isodepth captures the muscularis-lumen axis.

We next used GASTON-VI to identify proteins with differential spatial abundance gradients for the different pairs of anatomical regions of the intestine (Methods 4.5). We highlight two such proteins with significant differential gradients found by GASTON-VI. The immune marker CD161 has the largest DG score *D*_*m*_ and the smallest GASTON-VI *q* value when comparing spatial gradients between the duodenum and ileum (GASTON-VI *q* = 0.006, DG score *D*_*m*_ = 3.81 Fig. 5f). Interestingly, CD161 also exhibits differential spatial abundance gradients across other pairs of adjacent small intestine regions, including the duodenum versus proximal jejunum regions (GASTON-VI *q* = 0.006, DG score *D*_*m*_ = 0.72), the proximal versus mid jejunum regions (GASTON-VI *q* = 0.005, DG score *D* = 2.20), and the mid jejunum versus ileum regions (GASTON-VI *q* = 0.006, DG score *D* = 1.64). Visually, we observe that in the duodenum and proximal jejunum, CD161 abundance is largest near the lumen, while in the mid jejunum and ileum regions, CD161 abundance is largest near the muscularis (Fig. 5g). Because several immune cell populations express CD161, this change in spatial gradients across intestinal regions may reflect a shift in the distribution of the immune cells along the muscularis-lumen axis from proximal to distal small intestine. Another protein that GASTON-VI identified as having significant differential gradients is the immune marker CD4, which had the largest DG score between the ascending and transverse regions (GASTON-VI *q* = 0.007, DG score *D* = 3.12 Fig. 5h), which are adjacent regions in the colon. We also found that CD4 ranked among the top eight proteins by DG score between other adjacent colon regions including the transverse and descending colon, and descending and descending sigmoid colon, suggesting that CD4+ T cell enrichment along the muscularis-lumen axis may vary across different colonic segments (Fig. 5i). We note that our statistical test for differential spatial gradients was very efficient, taking only 19.1 minutes to run for a pair of conditions on a single NVIDIA L40S GPU.

## 3 Discussion

We introduce GASTON-VI, a deep learning model for learning an intrinsic spatial axis and spatial molecular gradients along this axis from large-scale, multi-sample and -condition spatial-omics data of any molecular modality. GASTON-VI is able to integrate multiple tissue slices from potentially different individuals, accounting for batch effects in order to recover a shared spatial axis and molecular gradients. Furthermore, GASTON-VI enables the identification of *differential* spatial gradients across different biological conditions. To the best of our knowledge, no existing computational methods are able to identify spatial axes from multiple tissue slices of different individuals and biological conditions nor perform differential spatial gradient analysis. At the same time, GASTON-VI is highly efficient to train, and thus is readily applicable to large-scale spatial-omics atlases with millions of cells.

We envision that the broad generality of our spatial gradient framework can be readily extended in numerous directions, including modeling interactions/correlations between molecular features (e.g. gene regulatory interactions, cell-cell communication) [84–90]; inferring spatial axes/gradients in tumors; inferring spatial gradients of RNA splicing parameters [93–97] and transcript diffusion; modeling spatial gradients in simulated ST data [100–102]; integration with large-scale foundation models [103–105]; and identifying sub-cellular molecular or morphological gradients [106–109]. Ultimately, the spatial axes and molecular gradients learned by GASTON-VI provide a versatile tool for characterizing spatial molecular variation in large-scale spatial-omics atlases.

## 4 Methods

### 4.1 A model of spatial and cell state variation in spatial-omics data

We derive a model of spatial and molecular variation along an underlying spatial axis in two-dimensional (2-D) tissue slices in spatial-omics data. We model each measured molecular feature (modality) *m* = 1, …, *M* in a 2-D tissue slice *T* ⊆ ℝ^2^ with a *molecular modality function f*_*m*_ : *T* × ℝ^*D*^ → ℝ, where *f*_*m*_(**s, w**) is the (normalized) abundance of molecular feature *m* in a cell at spatial location **s** = (*x, y*) ∈ *T* with cell state vector **w** = [*w*_*d*_] ∈ ℝ^*D*^. The cell state vector **w** describes intrinsic cellular factors such as cell type and cell cycle state. We assume the dimension *D* of the cell state vector **w** is known. For example, in spatial transcriptomics (ST) data, the function *f*_*m*_(**s, w**) is a *gene expression function* that gives the expression of gene *m* for a cell at location **s** with state **w**, while in spatial proteomics (SP) data, the function *f*_*m*_(**s, w**) is a *protein expression function* that gives the expression of protein *m* for a cell at location **s** with state **w**.

Estimating the molecular modality function *f*_*m*_ and cell state vector **w** is difficult due to the low sequence coverage (sparsity) and/or spatial resolution of current spatial-omics technologies. To avoid overfitting, we make the following assumptions on the molecular modality function *f*_*m*_.

First, we assume that the molecular modality function *f*_*m*_(**s, w**) is an *additively separable* function of the spatial coordinates **s** and cell state **w**:

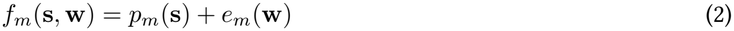

where *p*_*m*_ : *T* → ℝ and *e*_*m*_ : ℝ^*D*^ → ℝ are continuous functions called the *spatial modality function* and the *cell state modality function*, respectively. The spatial modality function *p*_*m*_(**s**) describes how the molecular feature *m* varies with a cell’s spatial location **s**, while the cell state modality function *e*_*m*_(**w**) describes how molecular feature *m* varies with the state **w** of a cell.

Second, we assume the spatial modality function *p*_*m*_ is a function of a *one-dimensional* (1-D) spatial coordinate *d*:

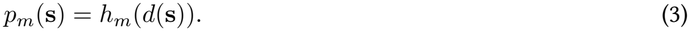

Here, *d* : *T* → ℝ is a continuous function called the *isodepth*, which describes the relative position of a spatial location **s** along an underlying 1-D spatial axis of the tissue slice *T*, and *h*_*m*_ : ℝ → ℝ is a continuous function which we call the *(1-D) spatial modality function*. We call *h*_*m*_(*d*(**s**)) a *1-D function* as it is a function of a single spatial coordinate *d*, while the (2-D) function *p*_*m*_(**s**) is a function of the 2-D spatial coordinates **s** = (*x, y*). The assumption in Eq. (3) reflects the observation that in many tissues, the molecular features that spatially vary will often vary continuously along an underlying 1-D spatial axis *d*; e.g. the PV-CV axis in the liver or the layer depth axis in the skin.

Under our two assumptions, the molecular modality function *f*_*m*_ has the form:

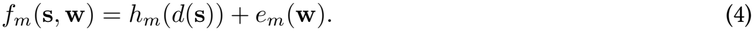

**Spatial gradients**. A key property of the isodepth *d* and the spatial modality function *h*_*m*_ are that they model *spatial gradients* of molecular feature abundance. Specifically, the spatial gradient ∇s*f*_*m*_(**s, w**) of molecular feature *m* is given by:

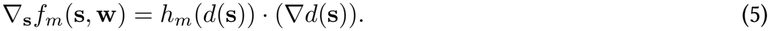

Thus, the spatial gradient ∇s*f*_*m*_ of each (spatially varying) molecular feature *m* is proportional to the spatial gradient ∇*d* of the isodepth, with proportionality constant given by the spatial modality function *h*_*m*_. Here, we will abuse notation and also refer to the spatial modality function *h*_*m*_ as a *spatial gradient*; although a spatial gradient is technically a vector field over physical space, it is common in biological literature to use the term spatial gradient to describe how a molecular feature varies along a 1-D spatial coordinate (e.g. [110–112]).

### 4.2 Extending to multiple tissue slices (batches) and conditions

We next extend our model to multiple tissue slices of the same underlying organ, which may be collected from different individuals with potentially different conditions (e.g. developmental or disease status). We are given *B* tissue slices *T*_1_, …, *T*_*B*_ ⊆ ℝ^2^, where we call each individual tissue slice *T*_*b*_ a *batch*. Each tissue slice *T*_*b*_ also has a discrete label *c*_*b*_ ∈ *C*, which we call a *condition*, for some finite set *C* of condition labels. For example, *c*_*b*_ ∈ {*H, D*} = *C* may indicate whether batch *b* comes from a healthy or diseased individual. (If there are no conditions labels specified, then we assume each tissue has a single condition *C*_*b*_ *= c* and the conditional label set |*C*| = 1 has a single element.)

We model spatial molecular variation in each batch *b* = *1*, …,*B* with a *batch-b molecular modality function f*_*mb*_ : ℝ^2^ × ℝ^D^ × *C* → ℝ, where *f*_*m*_*b(***s, w**, *c*) is the (normalized) value of molecular feature *m* in a cell at location **s** ∈ *T*_*b*_ with cell state w, where tissue slice *T*_*b*_ has condition c. Following Eq. (4), we assume that *f*_*mb*_ has the form

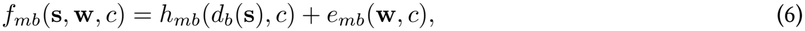

where *d*_*b*_ : *T*_*b*_ → ℝ is a continuous function which we call the *batch-b isodepth; h*_*mb*_ : ℝ × *C* → ℝ is a function, continuous in its first argument, which we call the *batch-b 1-D spatial modality function;* and *e*_*mb*_ : ℝ^*D*^× *C* → ℝ is a function, continuous in its first argument, which we call the *batch-b cell state modality value function*.

Since each batch *T*_*b*_ is a tissue slice from the same underlying tissue/organ (e.g. liver, intestine), the tissue slices share the same underlying isodepth axis and spatial molecular gradients along the isodepth axis. We model such shared structure by assuming that the batch-*b* 1-D spatial modality functions *h*_*mb*_ are *identical* across batches, i.e. *h*_*mb*_*(d, c)* = *h*_*m*_*(d, c)* for batches *b* = 1,. .., B for a given condition c. We note that the spatial modality functions *h*_*m*_*(d, c)* may be different across conditions c, reflecting that tissues from individuals in different conditions may have differential spatial gradients. We also note that each batch *T*_*b*_ may exhibit a different range(d_b_) of isodepth values, which may reflect factors including variation in resection of the tissue slice from the organ or variation of the underlying isodepth axis across biological conditions.

We also assume that the batch-*b* cell state modality function *e*_*mb*_ does not depend on the condition label *c*, i.e. *e*_*mb*_(**w**, *c)* = e_mb_(**w**). This assumption removes redundancy, as the batch *b* is already informative of the condition label *c*, and empirically improves inference of our model. Under our assumptions, each batch-*b* molecular modality function *f*_*mb*_ has the form

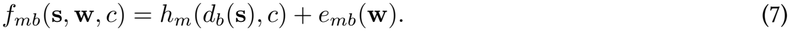

### 4.3 Generative modeling and the multi-batch spatial axis problem

We aim to estimate the batch-*b* isodepths *d*_*b*_, cell state vectors **w**_*ib*_, 1-D spatial modality functions *h*_*m*_, and the cell state modality functions *e*_*m*_ from the observed data. The observed data is multi-batch spatial-omics data {(**A**_*b*_, **S**_*b*_, *c*_*b*_*) }*_*b=1*,…,*B*_, where the b-th batch (tissue slice) (**A**_*b*_, **S**_*b*_, *c*_*b*_) consists of (1) a molecular modality matrix 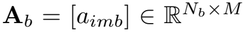, where each entry *a*_*imb*_ gives the value of molecular feature *m* = 1, .. ., M in cell *i* = *1*,…, *N*_*b*_; (2) a spatial coordinate matrix 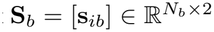 with entries **s**_*ib*_ = *(x*_*ib*_, *y*_*ib*_) ∈ ℝ^2^ giving the spatial coordinates of cell *i;* and (3) a condition label *c*_*b*_∈ *C*.

We formalize this problem using *generative modeling*, which involves defining a probability distribution for the observed data {(**A**_*b*_, **S**_*b*_, *c*_*b*_*) }*_*b=1*,…,*B*_ • We model the observed molecular features *a*_*imb*_ with a distribution ℙ (a_*imb*_ | *h*_*m*_, *e*_*m*_, *L*_*ib*_, Θ_*imb*_), where *h*_*m*_, *e*_*m*_ are the 1-D spatial and cell state modality functions in (7); Ƶ_*ib*_ are (unknown) *latent variables* for cell *i* in batch b; and Θ _m_ are (unknown) *modality-specific parameters*. The latent variables Ƶ_*ib*_ include the isodepth 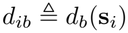 and state vector **w**_*ib*_ for the cell, as well as potential cell-specific confounders. The choice of the distribution depends on the specific modality being measured; here, we follow the standard approaches used in existing single cell generative models to model different molecular modalities (transcriptomics, proteomics) [36, 113], which we briefly describe below.

#### Spatial transcriptomics

We model the observed mRNA transcript count *a*_*imb*_ for gene *m* in cell *i* and batch *b* with a negative binomial (NB) distribution of the form:

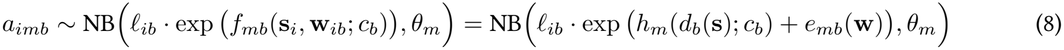

where *l*_*ib*_ *>* 0 is called the *library size factor* for cell *i* in batch *b* and *θ*_*m*_ *>* 0 is a gene-specific dispersion parameter. Each cell *i* in batch *b* has latent variables Ƶ_*ib*_ = *{d*_*ib*_, **w**_*ib*_, *l*_*ib*_*}*. We parametrize the *prior distributions* of these latent variables with the standard distributions used in single cell generative models [36,113], namely *d*_*ib*_ ∼ *N*(0,1), **w**_*ib*_ ∼ *N*(0, I), 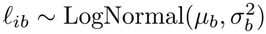, where μ_*b*_ and 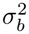 are the empirical mean and variance of the log-library size per batch *b* (see [36] for details). The modality-specific parameters are Θ_*m*_ = *{θ*_m_}.

### Spatial proteomics

We first process the raw protein intensities using an arcsinh transformation, following the DiagVI spatial-proteomics tutorial [114] (see Appendix S2.6). We model the processed intensity *a*_*imb*_ for protein *m* in cell *i* and batch *b* using a Gaussian likelihood, consistent with prior Gaussian modeling of single-cell proteomics data [114]:

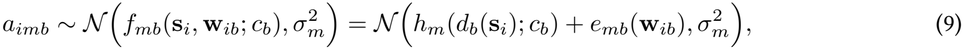

where *σ*_*m*_ *>* 0 is a protein-specific standard deviation term shared across cells *i* and batches *b*. Each cell *i* in batch *b* has latent variables *Ƶ*_*ib*_ = {*d*_*ib*_, **w**_*ib*_ *}*with Gaussian priors *d*_*ib*_ *∼N* (0, 1) and **w**_*ib*_ *∼ N* (0,*I*), and the modality-specic parameters are Θ _*m*_ = *{σ*_*m*_ *}* .

Given the generative model described above, we dene the multi-batch spatial axis problem (MSAP) as the following two problems: (1) estimate the parameters *h*_*m*_, *e*_*m*_, Θ _*m*_ that maximize the log-likelihood log *p*( {(**A**_*b*_, **S**_*b*_, *c*_*b*_) _*b*=1,…,*B*_ |*h*_*m*_, *e*_*m*_, Θ _*m*_) of the observed data {(**A**_*b*_, **S**_*b*_, *c*_*b*_)} _*b*=1,…,*B*_; and (2) compute the distribution *p*(*Ƶ* _*ib*_ |(**A**_*b*_, **S**_*b*_, *c*_*b*_), *h*_*m*_, *e*_*m*_, Θ_*m*_) (called the *posterior distribution*) of latent variables *Ƶ*_*ib*_ given the observed data (**A**_*b*_, **S**_*b*_, *c*_*b*_) and previously estimated parameters *h*_*m*_, *e*_*m*_, Θ _*m*_. Problem (1) yields the modality functions *h*_*m*_, *e*_*m*_, while problem (2) yields the (distribution of) the isodepth *d*_*b*_ and cell state vectors **w**_*ib*_. We formalize the MSAP below.

#### Multi-batch Spatial Axis Problem (MSAP)

*Given multi-batch spatial-omics data {*(**A**_*b*_, **S**_*b*_, *c*_*b*_) }_*b*=1,…,*B*_, *estimate the batch-b 1-D spatial modality functions h*_*m*_, *cell state modality functions e*_*m*_, *and modality-specic parameters* Θ_*m*_ *that maximize the log-likelihood of the data:*

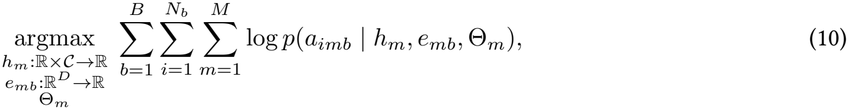

*and compute the distributions* 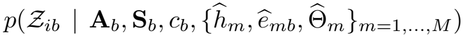 *where ĥ*_*m*_, *ê*_*mb*_, 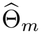 *are the solutions to*

### 4.4 GASTON-VI solves the MSAP with variational inference

The MSAP is a challenging problem as the objective of Equation (10) requires computing a computationally intractable integral, namely *p*(*a*_*imb*_| *h*_*m*_, *e*_*mb*_, Θ_*imb*_) = ∫*p*(*a*_*imb*_ *h*_*m*_, |*e*_*mb*_, Θ_*imb*_, *Ƶ*_*ib*_) *p*(*Ƶ* _*ib*_) *dƵ* _*ib*_. Here, we approximately solve the MSAP using *variational inference (VI)*, a statistical framework for approximating intractable integrals that is commonly used for multi-batch integration and analysis of single cell data (e.g. [36, 113]).

Briey, in VI, one maximizes a lower bound on the log-likelihood (i.e. the objective of (10)) called the *evidence lower bound objective (ELBO)*. The ELBO E is dened in terms of a distribution *q*_*ϕ*_ (*Ƶ*_*ib*_ | **A**_*b*_, **S**_*b*_) called the *variational posterior* that approximates the true posterior distribution; i.e. 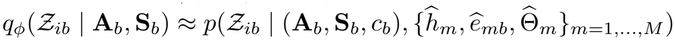 where *ϕ*are learned parameters. Here, we use a slight modication of the ELBO called the *(3*-weighted ELBO *ℰ*_*β*_[115] with additional regularization terms ℛ ( {*d*_*b*_}, *{***w**_*ib*_*}* ) for the isodepths *d*_*b*_ and cell states **w**_*ib*_. We specically solve the following optimization problem (see Appendix S1 for derivation):

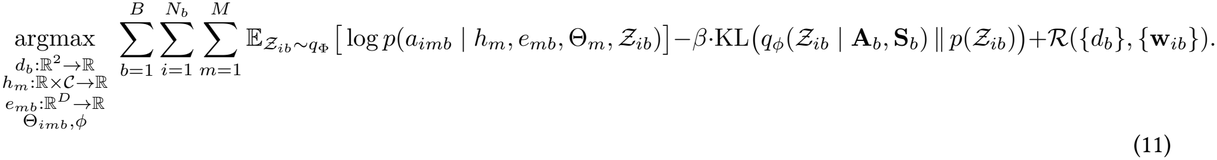

We implement the solution to (11) in an algorithm called GASTON-VI, with the following implementation choices.

#### Variational posterior

Following standard VI approaches [116], we parametrize the variational posterior distribution *q*_Φ_ (*Ƶ*_*ib*_ **a**_*ib*_, **s**_*ib*_, *b*) as the product of independent Gaussian distributions for the isodepth *d*_*ib*_ and cell state vector **w**_*ib*_; for transcriptomics data, we also use a log-normal distribution for the library size *l*_*ib*_ following [36]. See Appendix S1.1 for details.

#### Neural network architecture and training

GASTON-VI parametrizes the isodepth *d*_*b*_, cell state vector w, spatial modality function *h*_*m*_, and cell state modality function *e*_*mb*_ with multilayer perceptrons (MLPs). Spatial coordinates *s* are transformed into a higher-dimensional representation *γ* (*si*) using Fourier feature positional encoding [51]. We train GASTON-VI in two stages: an initial spatial warmup without the cell state w, followed by training of the full model. See Appendix S2.1–S2.3 for details.

#### Regularization terms *ℛ*

We use maximum mean discrepancy (MMD) regularization terms to reduce correlation between the isodepth *d* and cell state vector w and to encourage similar isodepth distributions across batches *b* with the same biological condition *cb*. See Appendix S2.4–S2.5 for details.

#### Modality-specific training

For CODEX spatial proteomics data, we additionally use an isodepth tail regularization term and rescale the inferred isodepth after each epoch for multi-batch training. See Appendix S2.6 for details.

#### Spatial gradient clustering

For each condition *c*, we use *K* -means clustering on the spatial modality functions *h*^*m*^*(d*, c) to identify *spatial gradient clusters ℳ*_*1*_,. .., *ℳ****K*** ⊆ {1,. .., *M*}, or groups of molecular features with similar spatial gradients *h*_*m*_*(d, c*). For each cluster *ℳ*, we call 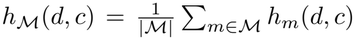 the cluster *ℳ*-averaged spatial modality function. See Appendix S2.7 for details.

### 4.5 A statistical test for differential spatial gradients

We next derive a statistical test to identify *differential* spatial gradients between two conditions *c*, c^’^, or whether the 1-D spatial modality functions *h*_*m*_*( ·c), h*_*m*_*(·,c’)* have the same shape or not. Without loss of generality (e.g. by scaling the isodepths), assume the range of isodepth values is given by *d*_*b*_*(x, y) 2* [0, 1] for all batches *b* in each condition. We define

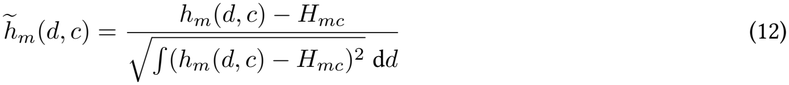

as the *normalized* 1-D spatial modality function for molecule *m* in condition c, where 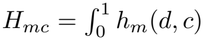 d*d* is the average of the 1-D spatial modality function *h*_*m*_*(· c)* across all isodepths *d*. The normalized 1-D spatial modality function 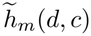 as mean 0 and variance 1 for *d 2* [0, 1].

For each molecule *m* and pair *c, c*^·^ of conditions, we aim to test between the following null and alternative hypotheses:

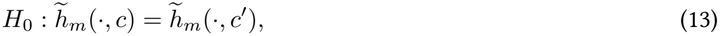

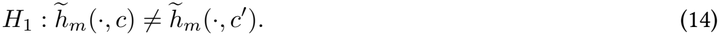

We note that we do not aim to test for direct equality of the 1-D spatial modality functions *h*_*m*_*(·, c), hm(·, c’*) since we only care about differences in the *shape* of the 1-D spatial modality function *h*_*m*_. For example, it is possible for a molecule *m* to have uniformly higher expression in one condition but for the spatial gradient curves *h*_*m*_*(d, c), h*_*m*_*(d, c’*) to have the same shape, e.g. if *h*_*m*_*(d, c’*) = *h*_*m*_*(d*, c) + *P* for some constant *P*.

We use the following test statistic:

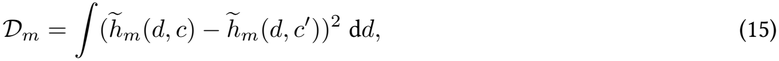

or the mean squared error between the normalized 1-D spatial modality functions *h*_*m*_*(d, c), h*_*m*_*(d, c’)*.

In practice, we approximate the test statistic *D*_*m*_ by clipping the inferred isodepth and discretizing the range of isodepth values to *T* discrete points *d*_1_ *<d*_2_ *<* ∆ *<d*_*T*_ and computing

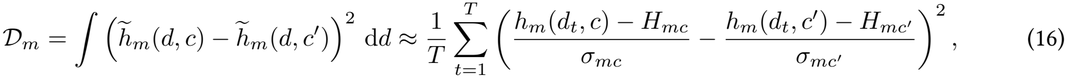

where 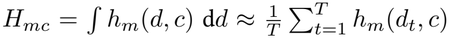 and 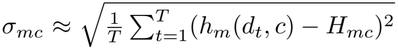 . We set d_1_ to be the δ-th quantile and *d*_*T*_ to be the (1 — δ)-quantile of overlapping isodepth values across the two conditions. In our implementation, we use *δ* = 0.02 and *T* = 60.

We estimate the null distribution of our test statistic *D*_*m*_ using a parametric bootstrap procedure. Briefly, we first refit a “null” GASTON-VI model by assigning all batches to the same condition, i.e. *c*_*b*_ = *c* for all batches *b*, so that each batch has the same 1-D spatial modality function *h*_*m*_(*d*). We then generate *B* synthetic molecular feature matrices **Ã**_*b*_ using the learned parameters from the null GASTON-VI model (e.g. for transcriptomics, sampling from the negative binomial in Eq. (8), or for proteomics, sampling from the Gaussian in Eq. (9)), yielding a synthetic multi-batch dataset *S* = {(**Ã**_*b*_, *S*_*b*_, *c*)}_*b*=1,…,*B*_ where each batch (**Ã**_*b*_, *S*_*b*_, *c*_*b*_) was generated given the same spatial modality functions *h*_*m*_(*d*). We repeat such a procedure *R* times to generate *R* different synthetic datasets *S*_1_, …, *S*_*R*_. We fit GASTON-VI to each synthetic dataset *S*_*r*_ using the original condition labels and compute the corresponding test statistic 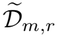, and we use the collection 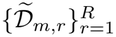 as our empirical null distribution for the test statistic *D*_*m*_. In our applications we use *R* = 200 null test statistics.

Using the estimated empirical null distribution 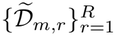, we calculate the empirical *p*-value for molecular feature *m* as

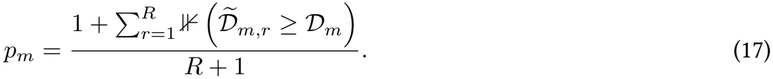

We adjust the resulting *p*-values across molecular features using the Benjamini-Hochberg procedure to control the false discovery rate, yielding adjusted *p*-values (*q*-values). We consider molecular features with *q* < 0.05 as having statistically significant differential gradients.

In practice, since refitting GASTON-VI *R* + 1 times may be computationally prohibitive, we improve the efficiency of our test by parallelizing the training of multiple GASTON-VI models on a single GPU concurrently.

### 4.6 Liver zonation metric

We quantified identification of known liver zonation genes using the area under the precision–recall curve (AUPRC). We computed AUPRC separately for each class in a one-vs-rest setting and reported the unweighted mean across classes (macro-averaged AUPRC; see sklearn documentation for details). To construct reference periportal and pericentral marker gene sets, we curated genes from existing literature that were either explicitly identified as canonical zonation markers in the literature [1, 60] or identified through statistical analysis in published datasets.

To compute the AUPRC for each method, for each gene *g* we fit a linear regression model between log-transformed gene expression values log(1 + *a*_*ig*_) and the latent coordinate *d*_*i*_ inferred by each model across spatial locations *i*, obtaining a slope *β*_*g*_ for each gene *g*. We rank genes according to their slopes; since either increasing or decreasing coordinates *d* may correspond to the PV-to-CV direction, we average the pericentral and periportal AUPRCs for both *d* and −*d* and report the higher mean AUPRC.

