## Supplementary Information for "GASTON-VI efficiently quantifies spatial axes and molecular gradients from multi-sample, multi-modal spatial-omics data"

### Appendix

#### S1 Variational Inference Formulation

To approximately solve the MSAP, we use VI to construct a tractable objective, the evidence lower bound (ELBO), which provides a lower bound on the marginal log-likelihood [54]. We maximize the ELBO as a tractable surrogate for maximizing the marginal log-likelihood in the MSAP (10).

To construct the ELBO, we approximate the intractable posterior over the latent variables isodepth $d_{ib}$, cell state vector $\mathbf{w}_{ib}$, and, for ST data, library size factor $\ell_{ib}$, with a variational posterior $q_\Phi$ with parameters $\Phi = \{\omega_1, \ldots, \omega_B, \phi, \tau\}$. We assume $q_\Phi$ to be factorized as follows:

$$q_\Phi(d_{ib}, \mathbf{w}_{ib}, \ell_{ib} \mid \mathbf{a}_{ib}, \mathbf{s}_{ib}, b) = q_{\omega_b}(d_{ib} \mid \mathbf{s}_{ib})\, q_\phi(\mathbf{w}_{ib} \mid \mathbf{a}_{ib}, b)\, q_\tau(\ell_{ib} \mid \mathbf{a}_{ib}, b). \tag{S1}$$

We jointly optimize neural-network parameters of the variational posterior $\Phi$, spatial molecular feature functions $h$, and cell-state molecular feature functions $e$, together with the inverse-dispersion parameters $\theta$. We define the $\beta$-weighted ELBO for cell $i$ in batch $b$ as

$$\begin{aligned} \mathrm{ELBO}^\beta_{ib}(h, e, \theta, \Phi) = \mathbb{E}_{q_\Phi}\big[\log p_{h,e,\theta}(\mathbf{a}_{ib} \mid d_{ib}, \mathbf{w}_{ib}, \ell_{ib}, b, c_b)\big] \\ - \beta\, \mathrm{KL}\big(q_{\omega_b}(d_{ib} \mid \mathbf{s}_{ib}) \,\|\, p(d_{ib})\big) \\ - \mathrm{KL}\big(q_\phi(\mathbf{w}_{ib} \mid \mathbf{a}_{ib}, b) \,\|\, p(\mathbf{w}_{ib})\big) \\ - \mathrm{KL}\big(q_\tau(\ell_{ib} \mid \mathbf{a}_{ib}, b) \,\|\, p(\ell_{ib} \mid b)\big), \end{aligned} \tag{S2}$$

where $p_{h,e,\theta}(\mathbf{a}_{ib} \mid d_{ib}, \mathbf{w}_{ib}, \ell_{ib}, b, c_b) = \prod_{m=1}^M \mathrm{NB}(a_{imb}; \ell_{ib}\rho_{imb}, \theta_m)$ is the NB likelihood for gene expression counts, with $\rho_{imb} = \exp(h_m(d_{ib}, c_b) + e_{m,b}(\mathbf{w}_{ib}))$. For other modalities, the likelihood may be replaced with an appropriate distribution, such as a Gaussian, and the library size factor $\ell_{ib}$ and its associated terms may be omitted. The Kullback–Leibler (KL) terms regularize the variational posteriors by penalizing deviations from their respective priors. Following $\beta$-VAE [115], we introduce a weight $\beta \geq 0$ on the isodepth KL term, with $\beta = 1$ recovering the standard VI ELBO. Since the isodepth prior is independent of spatial location, its KL penalty encourages a spatially constant isodepth function. We therefore empirically set $\beta = 0.01$ to reduce this penalty and allow the isodepth to capture complex spatial anatomical patterns.

Finally, we obtain a tractable approximation to the MSAP by maximizing the sum of the $\beta$-weighted ELBOs over all cells with respect to $h$, $e$, $\theta$, and $\Phi$:

$$\mathcal{L}(h, e, \theta, \Phi) = \sum_{b=1}^{B} \sum_{i=1}^{N_b} \mathrm{ELBO}^\beta_{ib}(h, e, \theta, \Phi). \tag{S3}$$

In practice, we augment $\mathcal{L}$ with additional regularization terms, described below.

##### S1.1 Variational posterior

Consistent with the priors, we model the variational posterior factors in (S1) for the isodepth and cell state with normal distributions and that of the library size factor with a log-normal distribution:

$$\begin{aligned} q_{\omega_b}(d_{ib} \mid \mathbf{s}_{ib}) &= \mathcal{N}\big(\mu_{\omega_b}(\mathbf{s}_{ib}),\, \sigma^2_{\omega_b}(\mathbf{s}_{ib})\big), \\ q_\phi(\mathbf{w}_{ib} \mid \mathbf{a}_{ib}, b) &= \mathcal{N}\big(\mu_\phi(\mathbf{a}_{ib}, b),\, \mathrm{diag}(\sigma^2_\phi(\mathbf{a}_{ib}, b))\big), \\ q_\tau(\ell_{ib} \mid \mathbf{a}_{ib}, b) &= \mathrm{LogNormal}\left(\mu_\tau(\mathbf{a}_{ib}, b),\, \sigma^2_\tau(\mathbf{a}_{ib}, b)\right), \end{aligned} \tag{S4}$$

where the distribution parameters are parametrized by neural networks with weights $\Phi$. We define the batch-$b$ isodepth $d_b$ in equation (7) as the posterior mean $d_b(\mathbf{s}) = \mu_{\omega_b}(\mathbf{s})$.

#### S2 GASTON-VI Implementation

We implement the VI algorithm described above in a software package which we call GASTON-VI. Below, we describe the neural network architecture and training procedures used to parameterize and optimize the model.

#### S2.1 Neural network architecture

GASTON-VI uses separate encoders for isodepth $d$, cell state latent $\mathbf{w}$, and library size $\ell$. Each batch-specific spatial encoder parameterizes the isodepth posterior $q_{\omega_b}(d_{ib} \mid \mathbf{s}_{ib})$ using a two-hidden-layer multilayer perceptron (MLP) with SiLU activations, followed by separate linear layers for the mean and standard deviation. We use 20 hidden units per layer for single-batch analyses and 64 for multi-batch analyses. The cell-state and library-size encoders parameterize $q_\phi(\mathbf{w}_{ib} \mid \mathbf{a}_{ib}, b)$ and $q_\tau(\ell_{ib} \mid \mathbf{a}_{ib}, b)$ following scVI [36], each with two hidden layers of 128 units and a dropout rate of $0.1$. For multi-batch analyses, both encoders include the batch label as a covariate.

The molecular feature function (Eq. (7)) is modeled using an additive decoder with separate spatial and cell state branches. The spatial branch uses a two-hidden-layer MLP with SiLU activations and 20 hidden units per layer to model the spatial modality function $h_m(d; c)$ from isodepth $d$ and an optional one-hot-encoded condition label $c$. The cell state branch uses a linear layer to model the cell state modality function $e_{mb}(\mathbf{w})$ from cell state $\mathbf{w}$ and an optional one-hot-encoded batch label $b$. The outputs of the two branches are summed to obtain the additive predictor $f_{m,b}$, which is transformed according to the modality-specific likelihood.

#### S2.2 Positional encoding

Standard coordinate-based MLPs exhibit a spectral bias toward low-frequency functions, limiting their ability to capture rapidly varying spatial anatomical patterns. This limitation is particularly relevant for tissues with repeated spatial organization, such as the liver, where PV-CV zonation recurs across lobules. To better capture these complex spatial anatomical patterns, we augment the spatial coordinates with Fourier feature positional encoding (PE) [51].

We standardize the spatial coordinates $\mathbf{s}_i = (x_i, y_i)$ using the training-set mean and standard deviation, with standardization performed separately within each batch in multi-batch analyses. We then map the standardized coordinates to a higher-dimensional sinusoidal representation:

$$\gamma(\mathbf{s}_i) = \left[\cos\left(2\pi\sigma^{j/\nu}x_i\right), \sin\left(2\pi\sigma^{j/\nu}x_i\right), \cos\left(2\pi\sigma^{j/\nu}y_i\right), \sin\left(2\pi\sigma^{j/\nu}y_i\right)\right]_{j=1}^{\nu}$$

where $\nu$ is the number of frequency bands and $\sigma$ controls the range of frequencies represented.

We use a data-driven heuristic to automatically select the PE hyperparameters $\nu$ and $\sigma$, with additional geometrically spaced wavelengths added when finer spatial scales are needed.

The final input to the spatial encoder is then $\gamma(\mathbf{s}_i) = \text{concat}\left(\mathbf{s}_i, \gamma(\mathbf{s}_i), \gamma_{\text{extra}}(\mathbf{s}_i)\right)$, where $\gamma_{\text{extra}}$ are the additional wavelengths if necessary.

#### S2.3 Two-Stage Training

We use a two-stage training procedure that first learns isodepth $d$ before optimizing the full model. During the first stage, or spatial warmup, we train a reduced model in which the cell state latent $\mathbf{w}$ is omitted, encouraging $d$ to capture the underlying 1-D spatial axis of molecular feature variation.

In the second stage, we then introduce the cell state latent $\mathbf{w}$ and optimize the full model, allowing $\mathbf{w}$ to capture residual sources of variation not explained by the isodepth. In all experiments, we use a spatial warmup of 100 epochs and train the full model for a total of 175 epochs.

#### S2.4 Disentanglement Regularization

To encourage the isodepth $d$ and cell state vector $\mathbf{w}$ to capture complementary sources of variation, we penalize their statistical dependence using a kernel-based maximum mean discrepancy (MMD) objective:

$$\mathcal{L}_{\text{MMD}} = \lambda\,\text{MMD}^2\big(q(d, \mathbf{w}),\ q(d)q(\mathbf{w})\big), \tag{S5}$$

where $q(d, \mathbf{w})$ denotes the aggregated joint posterior across cells, and $q(d)$ and $q(\mathbf{w})$ denote the marginal distributions. We use a Gaussian kernel with bandwidth 1 and set $\lambda = 0.01$.

After the spatial warmup phase, training alternates between optimizing the variational objective in Eq. (S3) and minimizing the disentanglement objective $\mathcal{L}_{\text{MMD}}$. We apply the MMD penalty only to the cell state encoder [28], keeping the learned isodepth fixed while encouraging the cell state encoder to remove spatial information from $\mathbf{w}_i$.

#### S2.5 Cross-batch isodepth alignment

In multi-batch analyses, GASTON-VI uses batch-specific spatial encoders and shared spatial modality function $h_m(d; c)$. To align the batch-specific isodepths $d_b$ to a common 1-D coordinate, we minimize the MMD between the isodepth distributions across all batch pairs with the same condition:

$$\mathcal{L}_{\text{align}} = \frac{\lambda_{\text{align}}}{|\mathcal{P}|} \sum_{(b,b')\in\mathcal{P}} \text{MMD}^2\left(\widehat{P}_b, \widehat{P}_{b'}\right), \tag{S6}$$

where $\widehat{P}_b$ and $\widehat{P}_{b'}$ denote the empirical distributions of inferred isodepth values (posterior mean) for batches $b$ and $b'$, respectively, and $\mathcal{P}$ is the set of batch pairs with the same condition $c$. $\lambda_{\text{align}}$ is set to 0.1.

#### S2.6 Training Modifications for CODEX Modality

For CODEX spatial proteomics data [15, 80], we replace the NB likelihood with a Gaussian likelihood. Protein features are first arcsinh-transformed (cofactor of 1), clipped at the 1st and 99th percentiles per protein, and min-max scaled to $[0, 1]$ within each anatomical region. Since CODEX measures protein intensities rather than sequencing counts, the library-size encoder is omitted and the decoder directly outputs the Gaussian mean without softmax normalization or scaling [114].

To prevent isodepth explosion during multi-batch training, we additionally apply an $\ell_2$ tail penalty,

$$\lambda_{\text{tail}}\,\text{ReLU}(|d| - \kappa)^2,$$

to isodepth values exceeding the cutoff $\kappa$. After each epoch, we rescale $d_\mu$ to zero mean and unit variance over the training set. All other architectural components and training procedures remain unchanged.

**Pre-processing.** We first apply an *arcsinh* transform with cofactor 1 to every protein value of every cell. *Arcsinh* transformation is commonly used for cytometry measurements and accommodates zero and negative values [119]. We then clip each protein's transformed values at the protein's 1st and 99th percentiles and rescale them to $[0, 1]$. The percentiles and the rescaling are computed separately for each anatomical region, using all cells from all batches corresponding to that anatomical region. *Arcsinh* transformation followed by feature-wise min-max scaling is also used in the DiagVI spatial-proteomics tutorial [114].

#### S2.7 Clustering spatial gene expression functions

To identify groups of genes with similar spatial expression patterns, we cluster the learned spatial expression functions $h_g(d, c)$ separately for each condition $c$ using K-means clustering. We refer to the average of the standardized spatial expression functions across all genes in a cluster $\mathcal{M}$ as the cluster-averaged spatial expression function, denoted by $h_{\mathcal{M}}(d, c)$.

We evaluate $h_g(d, c)$ on an isodepth grid $d_1, \dots, d_L$ and standardize each gene's $h_g(d, c)$ across the grid:

$$\widetilde{h}_g(d, c) = \frac{h_g(d, c) - \mu_{g,c}}{\sigma_{g,c}}, \tag{S7}$$

where $\mu_{g,c}$ and $\sigma_{g,c}$ are the mean and standard deviation of $h_g(d, c)$ over the grid. We apply K-means clustering to the vectors $(\widetilde{h}_g(d_1, c), \dots, \widetilde{h}_g(d_L, c))$. We evaluate $K = 2, \dots, 15$ and select $K$ using the elbow method based on K-means inertia.

For a gene cluster $\mathcal{M}$, we define its *cluster-averaged spatial expression function* as

$$h_{\mathcal{M}}(d, c) = \frac{1}{|\mathcal{M}|} \sum_{g\in\mathcal{M}} \widetilde{h}_g(d, c). \tag{S8}$$

For single-condition datasets, we omit the condition argument and write $h_g(d)$ and $h_{\mathcal{M}}(d)$.

### S3 GASTON-VI finds PV-CV axis and shared spatial gradients in healthy livers

We used GASTON-VI to analyze four MERFISH liver tissue slices collected from two healthy mice [58], profiling 347 genes across $1,275,831$ cells. Because all slices are sampled from healthy liver, we do not model condition-specific effects and learn only shared spatial gradients. GASTON-VI jointly models these tissue slices, or batches, by learning a batch-specific isodepth $d_b$ for each batch $b$ and spatial expression functions $h_g(d)$ that are *shared* across batches (Methods). GASTON-VI learns isodepths $d_b$ that recover the PV-CV axis in each tissue slice $b$, with isodepth values decreasing smoothly from the central vein towards the portal vein (Supplementary Fig. S2a). Jointly training GASTON-VI across all four tissue slices, which we call joint GASTON-VI, achieves comparable inference of the PV-CV axis and known liver zonation genes compared to training GASTON-VI individually on each slice, which we call individual GASTON-VIs, as quantified by AUPRC (Supplementary Fig. S2b). The shared spatial expression functions learned by joint GASTON-VI also recover the expected zonation gradients of known pericentral and periportal markers (Supplementary Fig. S2c), as well as the mid-zonal expression gradients of mid-zonal markers *Lyve1* and *Flt4*, whose gene expression peaks at intermediate positions along the PV-CV axis [1, 61] (Supplementary Fig. S2d). These results demonstrate that GASTON-VI accurately infers spatial organization and gradients shared across multiple tissue slices.

### S4 Learning cell state vector improves the spatial gradient found in liver

A key reason that GASTON-VI accurately infers spatial expression gradients along the PV-CV axis is because it also models cell state. Specifically, GASTON-VI models gene expression $f_g(\mathbf{s}, \mathbf{w})$ in terms of both the spatial location $\mathbf{s} = (x, y)$ *and* a learned cell state vector $\mathbf{w}$ that describes cell-intrinsic factors such as cell type. To demonstrate the importance of modeling cell state, we compared the 1-D spatial expression functions $h_g(d)$ learned by GASTON-VI and a variant of GASTON-VI without the cell state vector $\mathbf{w}$. We found that the spatial expression functions $h_g(d(\mathbf{s}))$ learned by GASTON-VI with the cell state vector $\mathbf{w}$ more accurately reflect the known spatial gradients of liver zonation genes compared to that learned by GASTON-VI without $\mathbf{w}$. For example, GASTON-VI with the cell state vector $\mathbf{w}$ learns a monotonically increasing 1-D expression function $h_g(d)$ for *Hsd17b12*, *Hsd17b2*, *Sardh*, and *Gpd1* (Supplementary Fig. S1), consistent with their status as known pericentral genes [2, 59, 61]. On the other hand, GASTON-VI without the cell state vector $\mathbf{w}$ learns spatial expression functions that do not reflect the known expression gradient for these pericentral genes (Supplementary Fig. S1). Our results demonstrate that explicitly modeling cell-intrinsic variation enables GASTON-VI to accurately characterize spatial gene expression gradients.

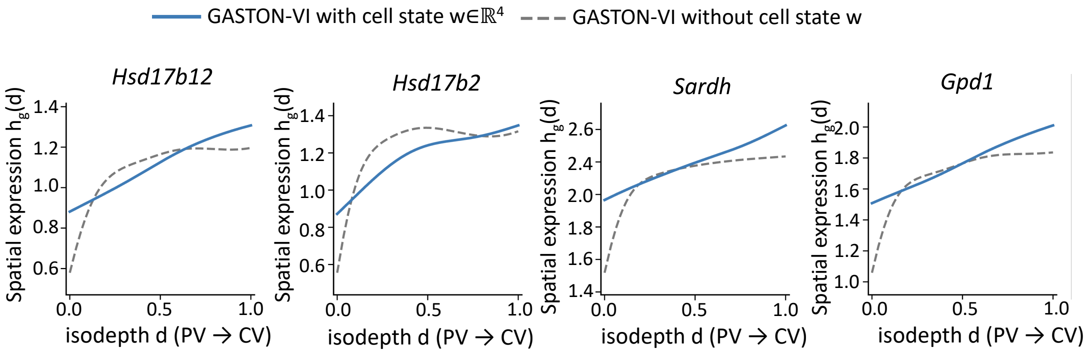


Figure S1: Spatial expression functions $h_g(d_b(\mathbf{s}))$ for representative pericentral markers *Hsd17b12*, *Hsd17b2*, *Sardh*, and *Gpd1* inferred by GASTON-VI with and without the cell state $\mathbf{w}$ on the MERFISH liver dataset [58].

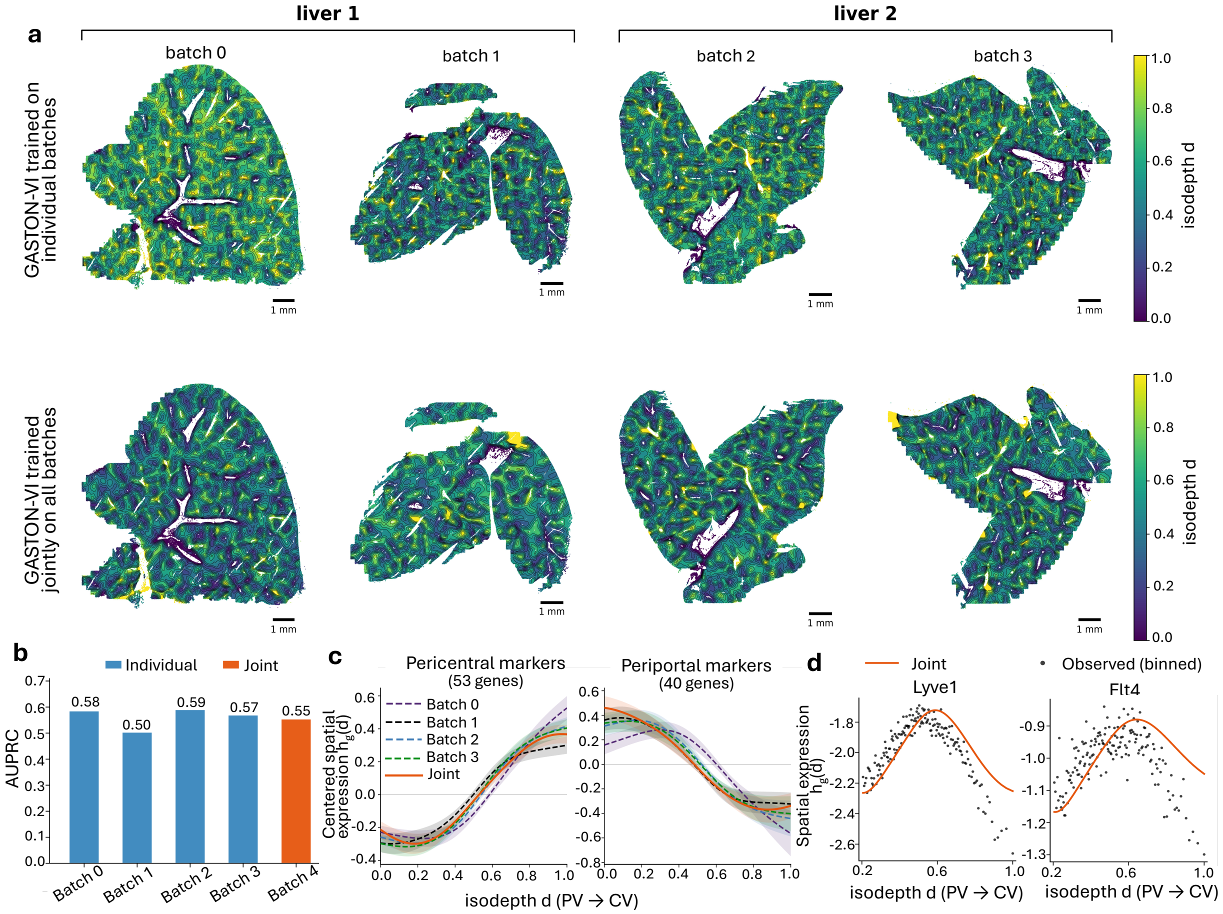


Figure S2: **a**, GASTON-VI isodepth learned on four MERFISH mouse liver tissue slices from two livers. Black curves denote contours of equal isodepth. Isodepth values were clipped to the 0.5th and 99.5th percentiles and globally min–max scaled across all four tissue slices for joint training. **b**, Comparison of zonation marker identification when GASTON-VI is trained individually on each batch versus jointly on all batches, quantified by AUPRC. **c**, Mean-centered 1-D spatial expression functions $h_g(d_b(\mathbf{s}))$ learned by GASTON-VI for pericentral and periportal marker genes when trained jointly on all batches versus individually. **d**, 1-D spatial expression functions $h_g(d_b(\mathbf{s}))$ learned by GASTON-VI for mid-zonated genes *Lyve1* and *Flt4* when trained jointly on all batches. Black dots denote the observed counts transformed to the same scale.

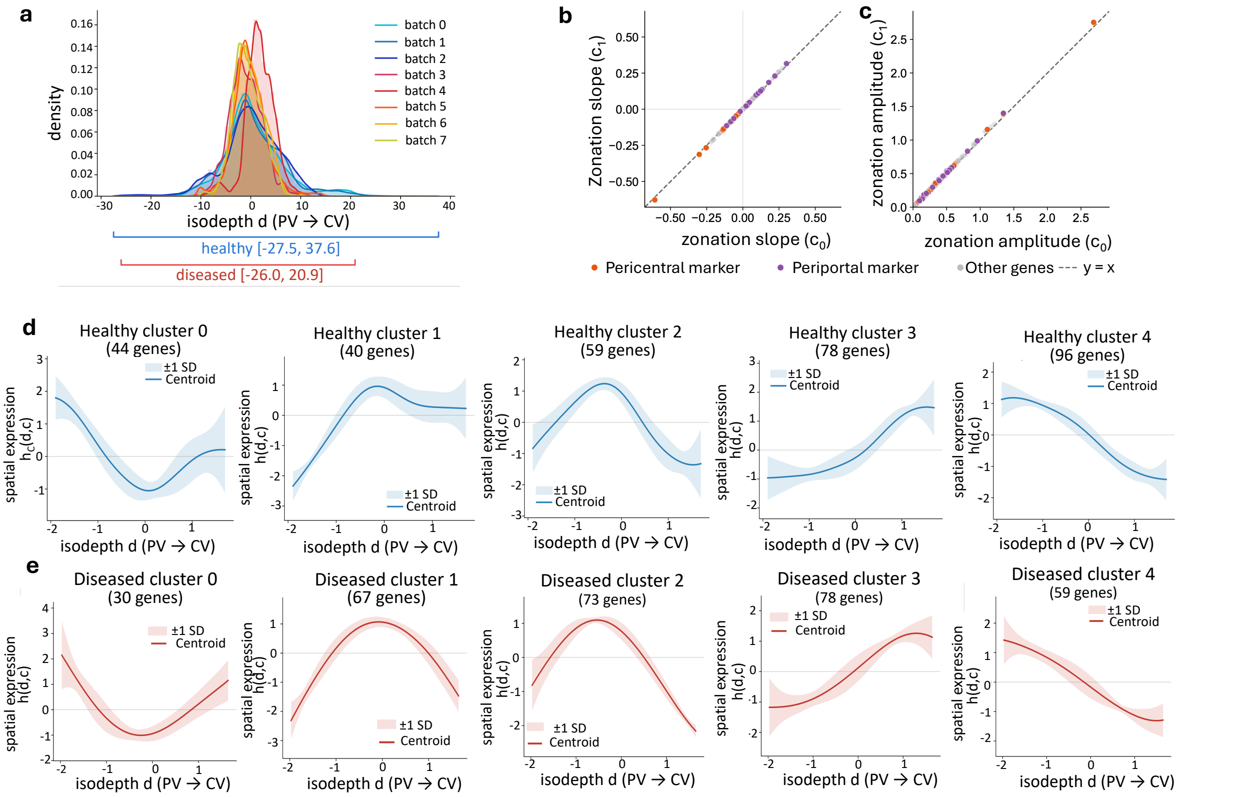


Figure S3: **a**, Distribution of the inferred isodepth for each batch. **b-c**, Negative control for condition-specific spatial expression patterns. Cells from healthy liver tissue were randomly assigned to one of two synthetic conditions, $c_0$ and $c_1$, within each batch. **b**, Comparison of zonation slopes between the two synthetic conditions. For each gene, the zonation slope was calculated by fitting a linear regression to the spatial expression function $h_g(d, c_0)$ and $h_g(d, c_1)$ against standardized isodepth $d$. Pearson $r > 0.999$, with a median absolute difference in slope of 0.003 between conditions. **c**, Comparison of zonation amplitudes between the two synthetic conditions. For each gene, zonation amplitude was calculated as the difference between the maximum and minimum values of $h_g(d, c)$ across isodepth. Pearson $r = 0.999$, with a median absolute difference in amplitude of 0.009 between conditions. **d**, Unsupervised clustering of spatial expression functions for healthy liver condition. Each gene's $h_g(d, c)$ was standardized across isodepth before clustering. The number of clusters was selected based on the Kneedle algorithm applied to the K-means inertia curve. **e**, Unsupervised clustering of spatial expression functions for diseased liver condition.

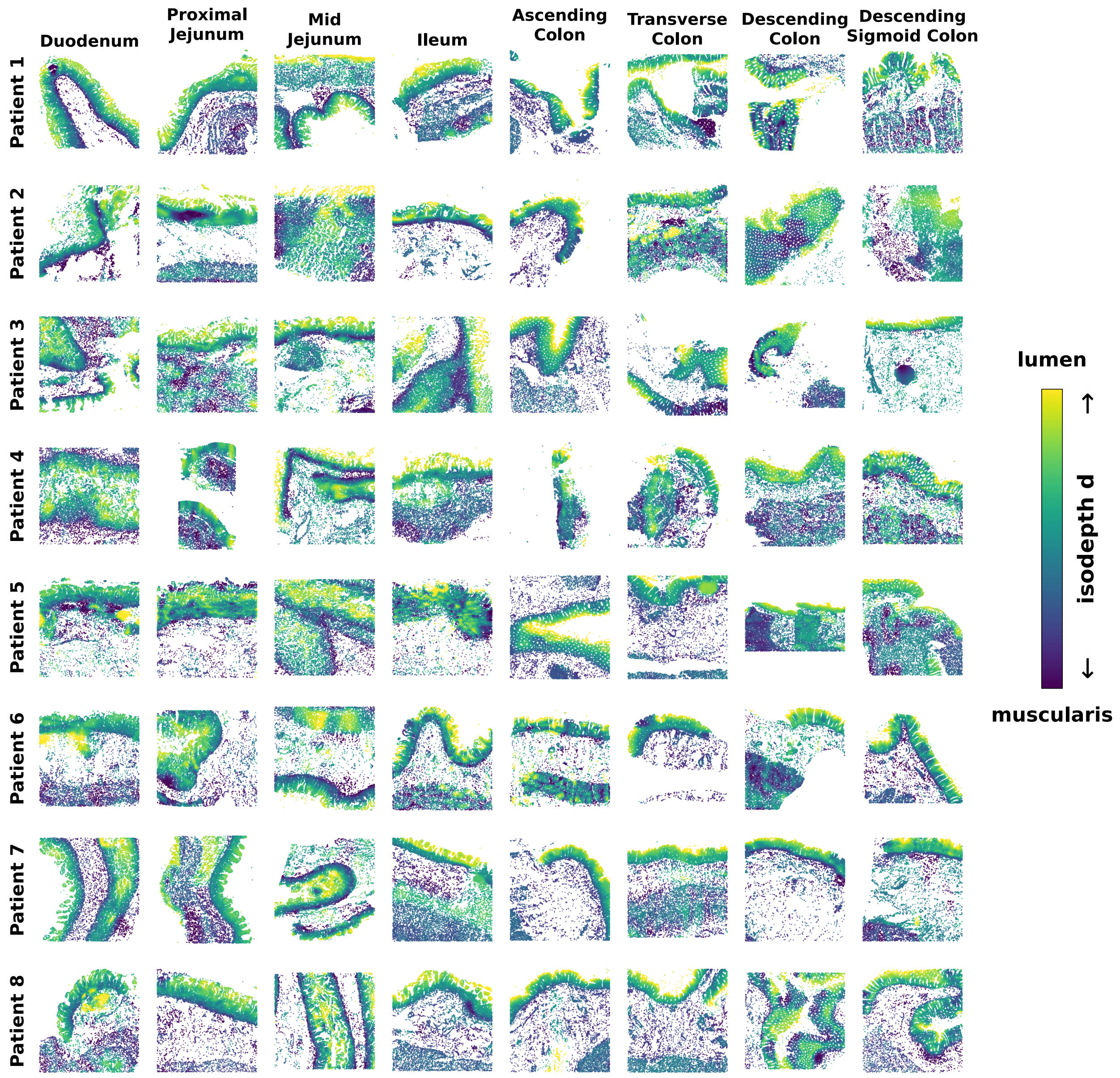


Figure S4: GASTON-VI isodepth learned across all 64 tissue slices from the human intestine CODEX spatial proteomics dataset [80]. Rows correspond to eight patients, and columns correspond to eight anatomical regions spanning the small intestine and colon. Each point represents a cell, colored by its inferred isodepth $d_b(\mathbf{s})$. Low isodepth values (purple) correspond to the muscularis side, and high values (yellow) correspond to the luminal side, revealing a shared muscularis-lumen axis across patients and anatomical regions.

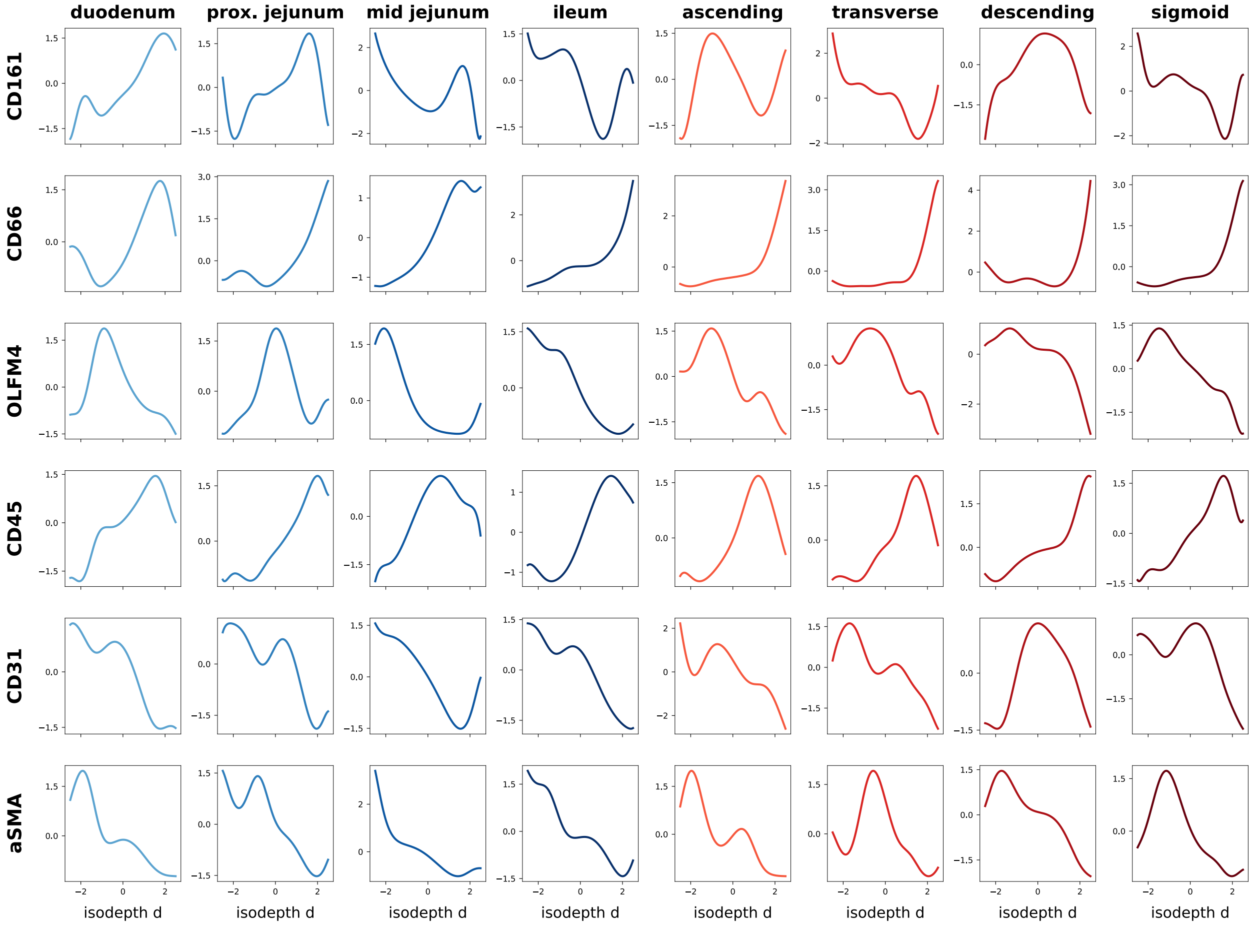


Figure S5: GASTON-VI spatial protein abundance functions $h_m(d, c)$ for six proteins (CD161, CD66, OLFM4, CD45, CD31, and $\alpha$SMA) across the eight intestinal conditions $c$ in the CODEX spatial proteomics dataset [80]. Rows correspond to proteins, and columns correspond to anatomical regions spanning the small intestine (blue) and colon (red). Each curve shows the learned spatial protein abundance function along isodepth $d$, with low values corresponding to the muscularis side and high values corresponding to the lumen side. CD161 (first row) exhibits differential spatial gradients, ranking third in the small intestine ($\mathrm{DG}_m = 0.83$) and eleventh in the colon ($\mathrm{DG}_m = 0.65$). CD66 ranks 40th in the small intestine ($\mathrm{DG}_m = 0.16$) and 48th in the colon ($\mathrm{DG}_m = 0.04$). OLFM4 ranks 16th in the small intestine ($\mathrm{DG}_m = 0.60$) and 47th in the colon ($\mathrm{DG}_m = 0.08$). CD45 ranks 41st in the small intestine ($\mathrm{DG}_m = 0.16$) and 46th in the colon ($\mathrm{DG}_m = 0.15$). CD31 ranks 49th (last) in the small intestine ($\mathrm{DG}_m = 0.09$) and 21st in the colon ($\mathrm{DG}_m = 0.45$). $\alpha$SMA ranks 45th in the small intestine ($\mathrm{DG}_m = 0.12$) and 33rd in the colon ($\mathrm{DG}_m = 0.30$).

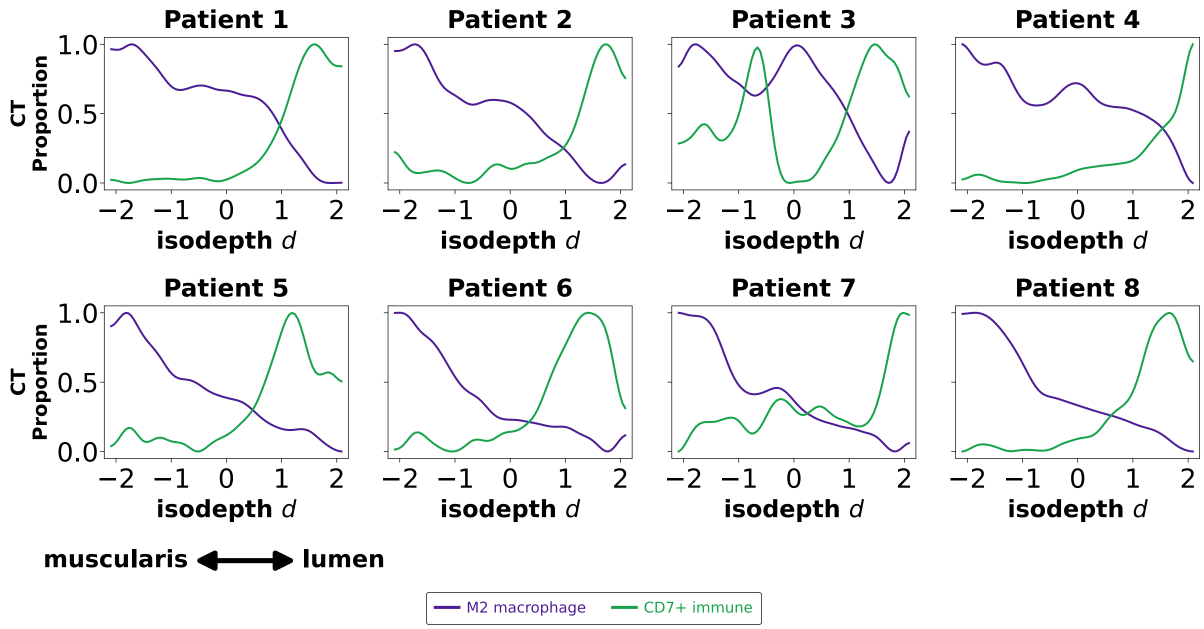


Figure S6: M2 macrophage and $CD7^+$ immune cell proportions along the GASTON-VI isodepth in the human intestine CODEX spatial proteomics dataset [80]. Each panel corresponds to one of eight patients and shows the proportions of M2 macrophages (purple) and $CD7^+$ immune cells (green) as functions of isodepth $d$. Low isodepth values correspond to the muscularis side, and high values correspond to the lumen side.

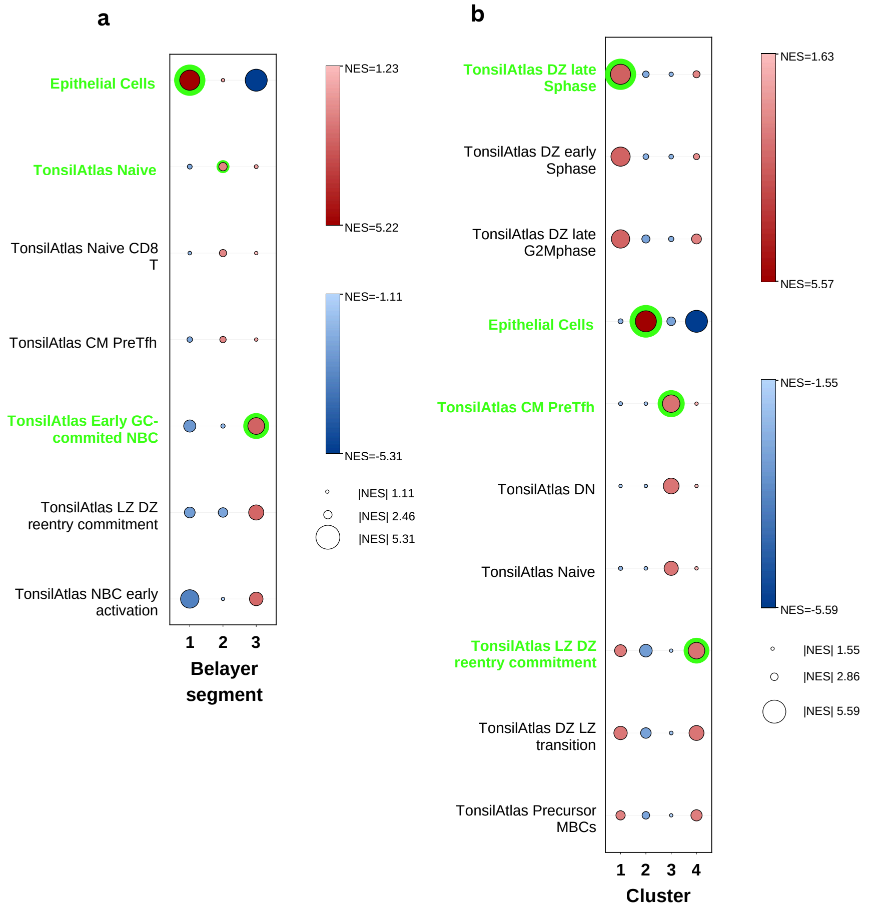


Figure S7: **GSEA validation of the tonsil isodepth axis.** For each of the three Belayer-segmented isodepth segments and each of the four unsupervised $h_{\mathcal{M}}(d)$ gene clusters, genes were ranked by expression/correlation and tested against a minimal reference library built from the Human Tonsil Cell Atlas and PanglaoDB. Dot color encodes normalized enrichment score (NES; red = positive, blue = negative) and dot size encodes $|\text{NES}|$; only FDR $< 0.05$ hits are shown. For each segment/cluster, the gene set with the highest NES value is outlined in green and its row label colored to match. **(a)** The three isodepth segments are enriched for an epithelial cell pathway (segment 1), a naive T cell pathway (segment 2), and a germinal-center B cell pathway (segment 3), validating the epithelium-TCZ-follicle axis along the isodepth $d$. **(b)** The four gene clusters are each enriched for multiple pathways, including a dark-zone B-cell late S-phase pathway (cluster 1), an epithelial cell pathway (cluster 2), a pre-Tfh cell pathway (cluster 3), and a light-zone to dark-zone reentry pathway (cluster 4). We therefore assigned broader cell-population labels to summarize their enrichment: effector B cells (cluster 1), epithelial cells (cluster 2), TCZ cells (cluster 3), and mixed immune/stromal cells (cluster 4).

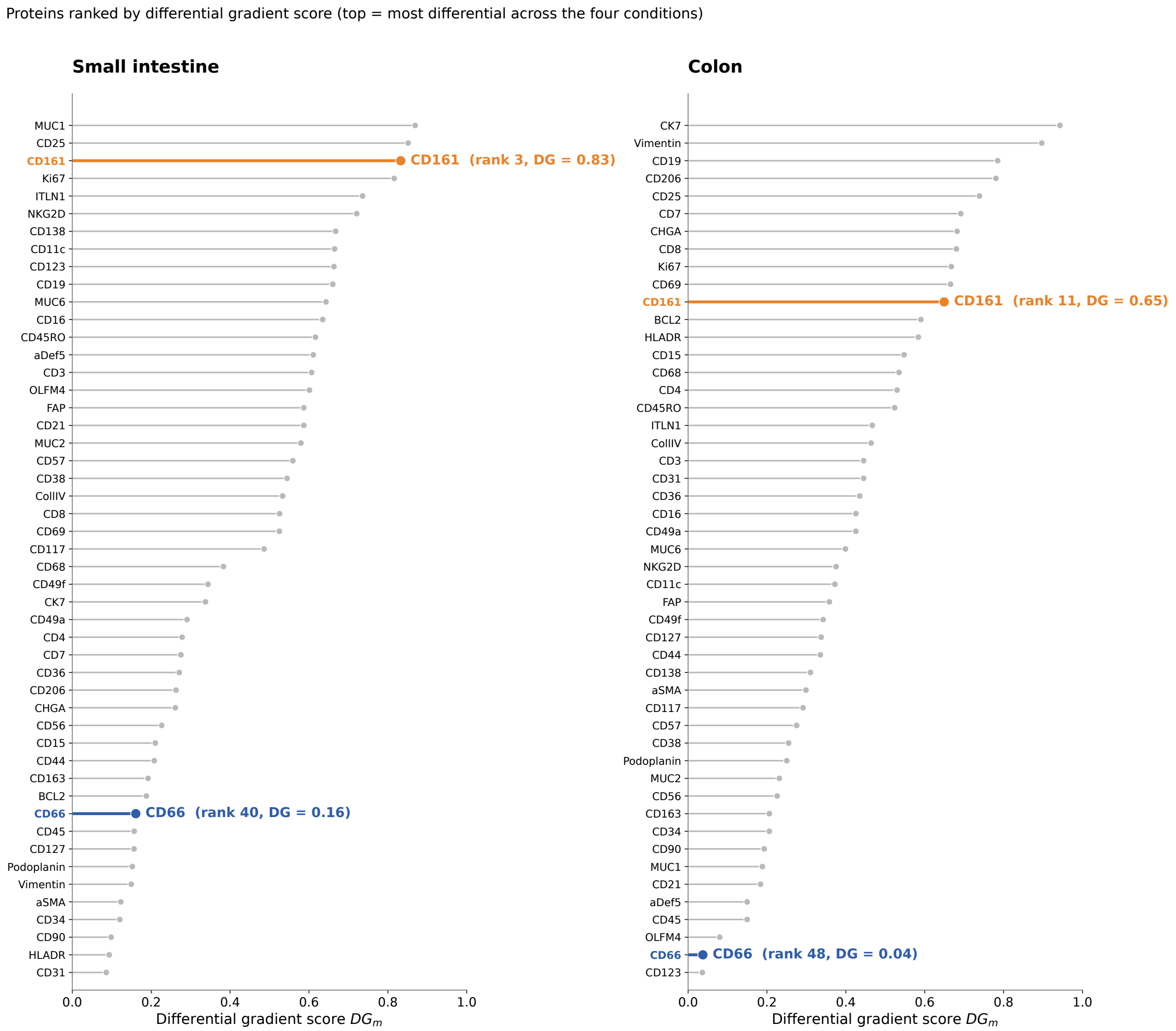


Figure S8: Ranking of all 49 proteins by differential gradient score $\mathrm{DG}_m$ in the human intestine CODEX spatial proteomics dataset [80]. Proteins are ranked from highest to lowest $\mathrm{DG}_m$ separately for the small intestine (left) and colon (right), based on differences in the shapes of spatial protein abundance functions $h_m(d, c)$ across the four anatomical conditions within each group. Higher scores indicate more differential spatial gradients across conditions, while lower scores indicate more shared gradients. CD161 ranks third in the small intestine ($\mathrm{DG}_m = 0.83$) and eleventh in the colon ($\mathrm{DG}_m = 0.65$), whereas CD66 ranks 40th in the small intestine ($\mathrm{DG}_m = 0.16$) and 48th in the colon ($\mathrm{DG}_m = 0.04$).
